# Sex-specific control of locomotor behavior by neuronal Arhgef10 in aged *Drosophila*

**DOI:** 10.64898/2025.12.18.694955

**Authors:** Mafalda Gualdino, Fabiana Herédia, Alexandra M. Medeiros, Rebeca Zanini, Juliane Menezes, Angela R. Mantas Dias, Márcia Garcez, Soren Prag, Antonio Jacinto, César S. Mendes, Andres Garelli, Alisson M. Gontijo

## Abstract

Mutations and polymorphisms in the *Rho guanine nucleotide exchange factor 10* (*ARHGEF10*) locus are associated with behavioral and locomotor dysfunctions in humans, including psychiatric disorders and polyneuropathies that show age- and sex-specific prevalence. *ARHGEF10* encodes a conserved guanine exchange factor that activates Rho GTPases and has proposed roles in cell migration and adhesion, but the relevant cell types and mechanisms underlying its age- and sex-dependent effects remain unclear. *Drosophila darhgef10* gene is the single orthologue of vertebrate *ARHGEF10* and its paralogue *ARHGEF10L*. Here, to gain more direct insight into possible age- and sex-dependent ARHGEF10 neuromuscular functions, we generated *darhgef10* knock-out flies and quantified induced walking kinematics with high-speed imaging and coarse spontaneous behaviors—walking, micromovements, rest, and sleep—in open arenas. Mutants of both sexes exhibited abnormal walking kinematics, which worsened with age in females. Cell-type-specific knockdowns indicated that locomotor phenotypes arose primarily from neuronal, rather than glial or muscle, requirements. dArhgef10 was especially critical in glutamatergic motor neurons of aged females. dArhgef10 was also required for wakefulness and activity initiation in females but, surprisingly, not in males. Genetic rescue and isoform expression analyses suggested that sexually-dimorphic expression of long isoforms RC and/or RD underlies at least part of the sex-specific requirements of dArhgef10 in locomotor behavior control, revealing unanticipated complexity in its activity regulation. Collectively, our results suggest ARHGEF10 plays an ancient, conserved role in neurons that promotes proper wakefulness and locomotor activity in an age- and sex-dependent manner.

## INTRODUCTION

Rho guanine-nucleotide exchange factors (ARHGEFs or GEFs) are proteins involved in the regulation of Rho family of small GTPases (Rho GTPases). Rho GTPases are a family of small G proteins belonging to the Ras superfamily of small GTPases. Human Rho GTPases are divided into 8 subfamilies that can be classified as typical and atypical depending on the way they are regulated. Typical Rho GTPase subfamilies include Rho, Rac/RhoG, Cdc42/RhoQ/RhoJ, and RhoF/RhoD. The Rho subfamily is composed by 3 members (RhoA-C), the Rac/RhoG subfamily by 4 members (Rac1-3 and RacG), the Cdc42/RhoQ/RhoJ subfamily by 3 members (Cdc42, RhoQ and RhoJ), and the RhoF/RhoD subfamily by 2 members (RhoF and RhoD) (Haga and Ridley, 2016; Heasman and Ridley, 2008). Whereas both typical and atypical Rho GTPases are activated when bound to Guanosine-5’-triphosphate (GTP), only the typical Rho GTPases cycle between an active (GTP-bound) and inactive (Guanosine-5’-diphosphate (GDP)-bound) state. The transition between active and inactive typical Rho GTPase (hereafter Rho GTPase) forms is most notably regulated by GEFs and GTPase-activating factors (GAPs): GEFs accelerate the exchange of Rho GTPase-bound GDP for GTP, leading to the activation of Rho GTPases, whereas GAPs promote GTP hydrolysis, resulting in Rho GTPase inactivation (Haga and Ridley, 2016; Heasman and Ridley, 2008; Hodge and Ridley, 2016). Once in active form, Rho GTPases can interact with effector proteins involved in the regulation of several cellular processes such as the regulation of gene expression, cell cycle, cytoskeleton remodeling, vesicle trafficking, cell morphogenesis, cell polarity, cell migration, cell division, and cell adhesion (Haga and Ridley, 2016; Heasman and Ridley, 2008; Hodge and Ridley, 2016). Well-studied examples of effector proteins of the Rho subfamily of Rho GTPase are the Rho-associated protein kinase (ROCK) and the Mammalian diaphanous-related Formin (mDia) (Haga and Ridley, 2016; Heasman and Ridley, 2008).

The human *ARHGEF10* gene codifies the Rho guanine nucleotide exchange factor 10 (ARHGEF10) protein of the Diffuse B-cell Lymphoma (DBL) family (Verhoeven *et al.,* 2003). DBL family members have a catalytic Dbl-homology (DH) domain through which they perform their GEF activity, and a pleckstrin homology (PH) domain thought to play a role in targeting RhoGEFs to plasma membrane and/or in assisting GEF activity (Haga and Ridley, 2016; Heasman and Ridley, 2008; Hodge and Ridley, 2016). Since the ARHGEF10 PH domain diverges from the PH domain typically found in the other DBL family GEFs, it has been considered a member of a subset of GEFs with a divergent or absent PH domain (Mohl *et al.,* 2006; Winkler *et al.,* 2005). This subset of GEFs also includes the ARHGEF10 paralogue, ARHGEF10L (also known as Grinch GEF), which also has a divergent PH domain, and ARHGEF17 (also known as p164-RhoGEF), which lacks this domain (Mohl *et al.,* 2006; Winkler *et al.,* 2005; Rümenapp *et al.,* 2002). All three members of this subset of GEFs also contain an additional carboxy (C)-terminal WD40 domain, which is thought to play a role in mediating the GEF interaction with other proteins (Mohl *et al.,* 2006; Jain and Pandey, 2018). *ARHGEF10* and *ARHGEF10L* are represented in *Drosophila* by a single gene, the *ARHGEF10/L* orthologue *CG43658*, which we refer hereafter to as *darhgef10* (**Figure 1A**). The *Drosophila ARHGEF17* orthologue is encoded by the gene *CG43102*.

**Figure 1.**
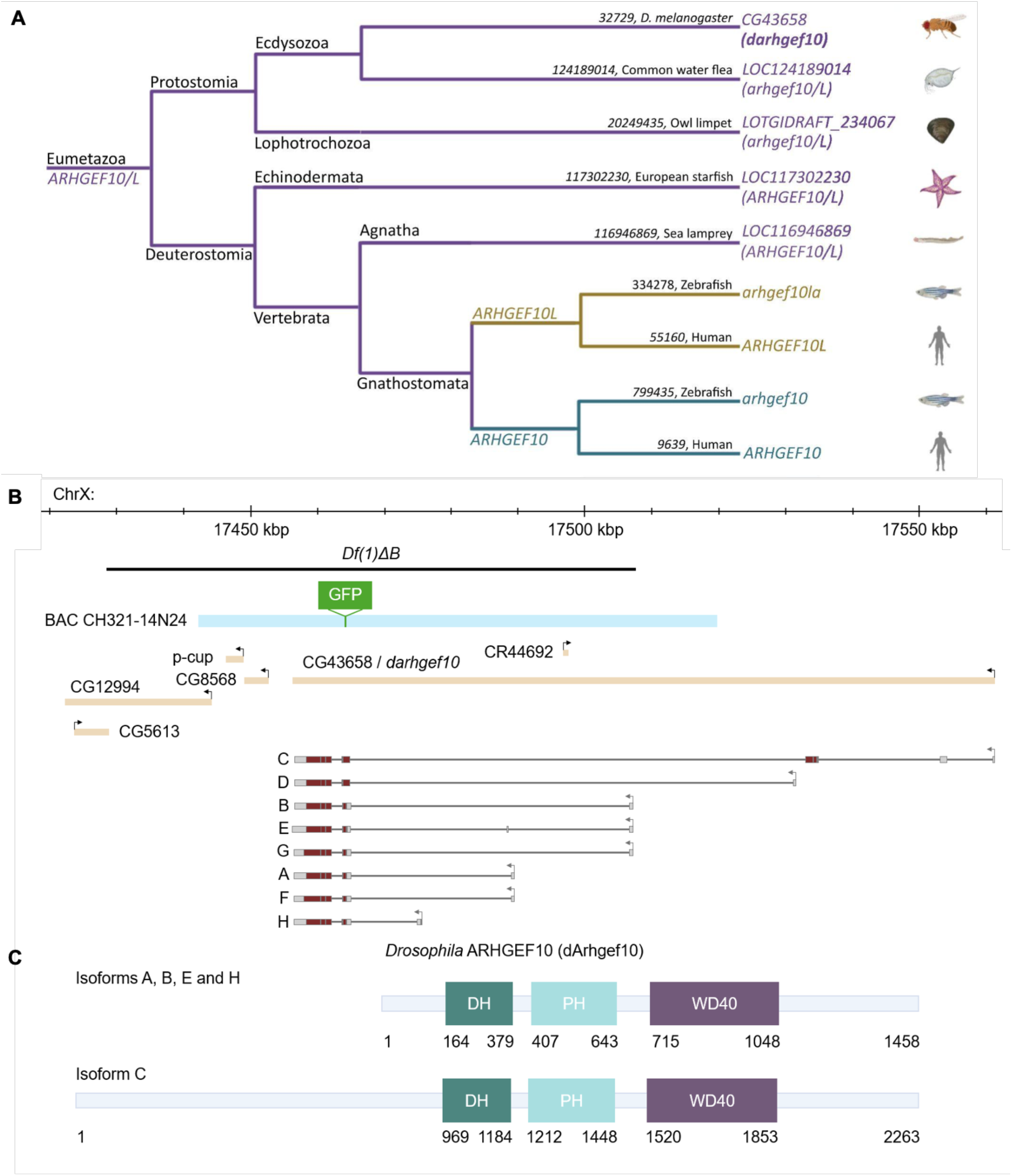
*ARHGEF10/ARHGEF10L* family relationships in metazoans and the *Drosophila melanogaster dahrgef10* locus and its protein isoforms. **(A)** Tree depicting the relationship between *ARHGEF10* family members in metazoans. Depicted are GeneID numbers (black) and official gene names (right) for known and predicted *ARHGEF10* and *ARHGEF10L* genes, or their predicted common ancestor or sole representative in a particular genome (*ARHGEF10/L*), which are generally most similar to *ARHGEF10* than to *ARHGEF10L*, hence the reason why the *Drosophila* orthologue is herein named *darhgef10* and not *darhgef10/L*. **(B)** Representative scheme of the *darhgef10* locus, showing the region deleted in *darhgef10* mutant flies, and the protein structure. The image depicts the *darhgef10* gene locus on the X chromosome and the deletion (*darhgef10* deficiency 1, or delta B for “big deletion” *[Df(1)ΔB*]) (black line, top). The image also depicts the eight known *darhgef10* transcript isoforms (A-H), and the five neighboring genes (*CG8568*, *p-cup*, *CG12994*, *CG5613* and *CR44692*) affected by *Df(1)ΔB*. The light blue line at the top of the scheme represents the *darhgef10* region included in the *darhgef10* rescue construct, which consists of a carboxy (C)-terminally GFP-tagged dArhgef10 (GFP-dArhgef10, also designated BAC CH321-14N24). Note that the “rescue” construct does not include the promoters and other elements of isoforms C and D. Hence, the rescue construct is not expected to fully recapitulate endogenous *darhgef10* expression. (C) *Drosophila melanogaste*r ARHGEF10 (dArhgef10) and its principal domains: Dbl homology (DH) (brown), Pleckstrin homology (PH) (gray), and WD40 (light brown). Adapted from interpro.

Mammalian ARHGEF10 has been shown to activate Rho GTPases of the Rho subfamily, which includes RhoA, RhoB, and RhoC (Mohl *et al.,* 2006; Chaya *et al.,* 2011), and there is evidence indicating that ARHGEF10 is involved in the regulation of cell contractility in a Rho-Rho-associated protein kinase (ROCK) dependent manner (Chaya *et al.,* 2011). It was also reported that ARHGEF10 localizes to cytoplasmic vesicles that move along microtubules regulating membrane trafficking, and that it is involved in the regulation of both cell migration (Shibata *et al.,* 2016; Joseph *et al.,* 2020), and cell-cell tight junctions (Khan *et al.,* 2021).

ARHGEF10 has also been shown to be critical for proper peripheral nervous system function (Verhoeven *et al.,* 2003; Høyer *et al.,* 2014). A mutation–consisting in the substitution of threonine for isoleucine at position 332 (Thr332Ile)–in the amino (N)-terminal region of ARHGEF10, upstream to the DH domain–was identified in humans with slow nerve conduction velocity associated with thin myelin fibers, but without clinical symptoms of neuropathy (Verhoeven *et al.,* 2003). Another mutation consisting in the substitution of arginine for threonine at position 338 (Arg338Thr), close to the previously-described Thr332Ile mutation, was identified in a patient with Charcot-Marie-Tooth (CMT) disease, a hereditary neuropathy characterized by slow nerve conduction, muscle weakness, and walking difficulties (Høyer *et al.,* 2014). Interestingly, a cell culture study showed that overexpression of either ARHGEF10 with its N-terminal region deleted [1-332 amino acid (aa) deletion], or of ARHGEF10 mutated at Thr332Ile, led not only to an increase in the amount of activated Rho GTPases of the Rho subfamily (RhoA, RhoB, and RhoC), but also to induced cell contraction compared to cells that overexpress normal ARHGEF10 (Chaya *et al.,* 2011). These findings suggest that the slow nerve conduction velocity associated with thin myelin fibers observed in the patients described above could be caused by increased RhoA-dependent cell contractility due to enhanced ARHGEF10 GEF activity. Furthermore, an ARHGEF10 coding region variant (*rs9657362*), which consists of a nonsynonymous substitution expected to lead to a leucine to phenylalanine change at position 370 (Leu370Phe) of ARHGEF10 has been associated with protection from chemotherapy-induced peripheral neuropathy in human patients treated with Taxol-derived compounds, which are drugs that affect microtubule function during cell division (Beutler *et al.,* 2014; Boora *et al.,* 2015). Finally, a homozygous deletion of 10 bp found in some large dog breeds is highly associated with an inherited polyneuropathy (Ekenstedt *et al.,* 2014), which shares clinical similarities to the Charcot-Marie-Tooth disease observed in humans. The 10 bp deletion detected in the canine *ARHGEF10* gene occurs within the exon 17-intron 17 junction and consists of the deletion of four nucleotides from the 3’-end of exon 17 and six nucleotides from the 5’-end of intron 17. This alteration is responsible for the generation of a frameshift in the processed messenger RNA (mRNA) which leads to the formation of a premature stop codon predicted to truncate approximately 50% of the protein (Ekenstedt *et al.,* 2014).

ARHGEF10 is also linked to central neurodevelopmental disorders associated with locomotor and behavioral dysfunctions in several organisms, including humans (Catusi *et al.,* 2021; Jungerius *et al.,* 2008; Lu *et al.,* 2018; Zhang *et al.,* 2022). In humans, various deletions in the 8p23.3 region affecting *ARHGEF10* and/or several neighboring genes including *FBXO25*, *DLGAP2*, *CLN8*, and *MYOM2*, are associated with developmental delay, motor coordination problems, attention deficit, hyperactivity, aggressiveness, and epilepsy (Catusi *et al.,* 2021). The deletion that only affects the *arhgef10* locus consists of the deletion of aa 1-12. Moreover, an association between the *ARHGEF10* intronic SNP, *rs11136442*, and schizophrenia development risk was observed in the Dutch population (Jungerius *et al.,* 2008). An Autism Spectrum Disorder (ASD) phenotype, characterized by social impairment, and hyperactivity, was observed in *arhgef10* knockout mice (Lu *et al.,* 2018), and locomotor abnormalities were observed in zebrafish larvae (Zhang *et al.,* 2022). The *arhgef10* deletion present in zebrafish larvae consists of the deletion of 121 bp and an insertion of 16 bp, which generates a frameshift mutation, and the production of a truncated protein with all the known functional domains disrupted (Zhang *et al.,* 2022).

Importantly, some diseases associated with mutations in *ARHGEF10* show age and sex differences. In schizophrenia, there are sex-specific differences in age of disease onset, disease outcome, and treatment (Li *et al.,* 2016). In men, schizophrenia onset occurs at an earlier age compared with women. Men have more severe negative symptoms like aggressiveness, social withdrawal, and substance abuse, while women present more mood disturbance, anxiety, depression, and affective symptoms. Finally, schizophrenic patients show sex differences in both antipsychotic drug dosage requirements and treatment response. While men require higher doses of antipsychotic drugs, women show better treatment compliance and response, and more side effects than men (Li *et al.,* 2016). Autism Spectrum Disorder (ASD) is more prevalent in male than in female patients and studies suggest sex differences in clinical manifestations, which, however, are still inconsistent because of difficulty in diagnosis (Napolitano *et al.,* 2022). In peripheral polyneuropathies, sex differences regarding age disease onset and prevalence were also documented. For instance, the *ARHGEF10* homozygous mutation found in some large breed dogs is related to juvenile-onset peripheral polyneuropathy, which is more prevalent in males. However, disease onset occurs in aged heterozygous or homozygous normal *ARHGEF10* animals (Ekenstedt *et al.,* 2014). In humans, peripheral neuropathies are more prevalent in the elderly and females, and 70-80% of the cases are associated with risk factors such as diabetes, alcohol overconsumption, cytostatic drugs, and cardiovascular disease (Hanewinckel *et al.,* 2016).

The molecular and cellular mechanisms associated with mutations in *ARHGEF10* are still unclear, and it is not known whether the processes in which ARHGEF10 is associated with are directly related to locomotor behavior control. Here, to gain insight into ARHGEF10 function in neuromuscular development and function, we performed loss-of-function studies of *darhgef10* in *Drosophila* using classical genetics and tissue-specific RNA interference (RNAi) and characterized fly locomotor function using different assays. Our results show that *darhgef10* is critical for proper *Drosophila* locomotor behavior, suggesting that *ARHGEF10* has an ancient, conserved role in controlling locomotor function, and further indicates that dArhgef10 is mostly required in neurons for proper locomotion in adults. Finally, our work reveals unexpected sex-specific requirements for dArhgef10 in *Drosophila.* Genetic rescue and isoform expression analyses suggested that long *darhgef10* isoforms RC and/or RD appear largely dispensable for proper spontaneous behaviors in females, but are transcriptionally enriched and functionally critical for proper behavioral regulation in males, likely acting to balance the activity of the other isoforms. These results reveal that sexually-dimorphic isoform expression can underly at least part of the sex-specific requirements of dArhgef10 in locomotor behavior control.

## RESULTS

### *darhgef10* mutant flies have an abnormal walking kinematics that worsens with age

To study *darhgef10* function in *Drosophila*, we generated a large deletion, *darhgef10* deficiency 1, or delta B for “big deletion” (*darhgef10[Df(1)ΔB]*), using FRT-mediated recombination (Parks *et al.,* 2004) (**Supplementary Figure S1**). *darhgef10[Df(1)ΔB]* completely deletes the common regions coding for the DH and PH-like domains in all isoforms (**Figure 1B**). No genomic sequence or transcripts are detected containing these domains, thus, we can predict that *darhgef10[Df(1)ΔB]* is a protein null (**Figure 1C and Supplementary Figure S1**). *darhgef10[Df(1)ΔB]* also completely or partially deletes four genes lying 5’ of *darhgef10* (*CG8568*, *p-cup*, *CG12994*, and *CG5613*), and a gene located in the + strand which codifies a long non-coding RNA (*CR44692*) (**Figure 1B**). *darhgef10[Df(1)ΔB]* female and male mutants–hereafter named, *darhgef10[-/-]* and *darhgef10[-]/Y*–are both viable and fertile.

As some of the disorders linked to *ARHGEF10* mutations are age-dependent, we first profiled the locomotor behavior of control and *darhgef10* mutants in young (2-day old) and aged (20-day old) female flies using the flywalker system (Mendes *et al.,* 2013). This assay uses an optical method (frustrated Total Internal Reflection) coupled with high-speed video imaging and dedicated software to automatically track the footprints and the body of walking flies. It enables the quantification of a large number of locomotor parameters–namely, step, spatial, gait, and stability parameters–that can be used to describe induced locomotor behavior with high temporal and spatial resolution (see **Supplementary Table S.2** for a full description of each locomotor parameter) (Mendes *et al.,* 2013).

At day 2-post-eclosion, a total of 16 out of 50 locomotor parameters across different kinematic categories changed significantly in young *darhgef10[-/-]* females relative to young control females (**Figure 2A, left column**). These changes, which were all relatively minor in effect size (typically <0.5 Log2 Fold change), included, for instance, slightly decreased speed and slightly increased stance straightness and non-canonical index. These results indicate that dArhgef10 plays a detectable, albeit small, role in promoting normal locomotion in young female flies.

**Figure 2.**
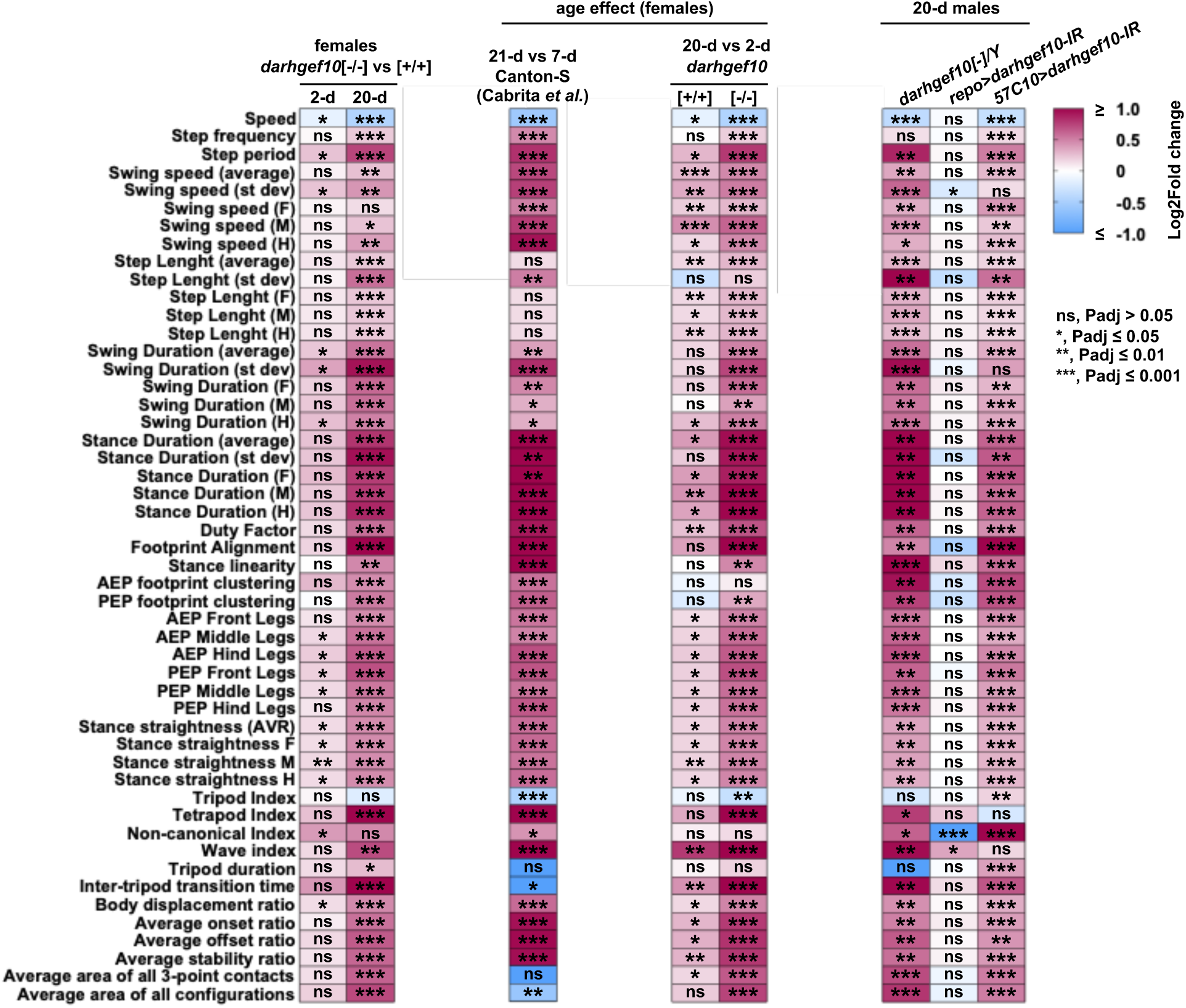
Neuronal dArhgef10 mitigates the effects of aging on locomotor behavior in female and male flies. **(A)** The effect of *arhgef10* on walking kinematics in females. Heatmap of the kinematic parameters of *darhgef10* mutant females (*darhgef10[-/-]*) aged 2 (left) and 20 (right) days after eclosion. For each parameter values are compared to age-matched control animals (“WT”, *darhgef10[+/+]*). **(B)** The effect of age on female walking kinematics. Heatmap of the kinematic parameters of aged vs young female flies: 3 (21-d)-vs. 1 (7-d)-week old WT control females (of the Canton-S genetic background) (data reanalyzed and replotted from the Cabrita *et al*. 2020) (left panel). 20-vs. 2-day old *darhgef10[+/+]* WT control (middle panel) and *darhgef10[-/-]* mutant females (right panel). **(C)** Pan-neuronal *darhgef10* knockdown phenocopies walking kinematics defects of aged male *darhgef10* mutants. Heatmap of the kinematic parameters of aged male *darhgef10* mutants (*darhgef10[-]/Y*) (left column), pan-glial knockdown (*repo>darhgef10-IR*) (middle column), and pan-neuronal knockdown (*57C10>darhgef10-IR*) (right column) males. For each parameter, values are matched to same age control animals. The effect size (Log2 Fold change) values are represented by a color code with red and blue shades indicating an increase or decrease relative to control, respectively. White indicates no variation. (n = 20 for each condition). Statistical significance was determined by two-tailed t-test: *P* ≤ 0.05 (\**), P* ≤ 0.01 (**), *P ≤ 0.001 (\*\*\**), ns = not significant (*P* > 0.05).

To study the kinematics of aged *darhgef10[-/-]* females, we first asked what was the effect of age alone on these parameters by comparing 2-vs. 20-day old control female flies. We found that age alone had a drastic effect on locomotor parameters, affecting 34 out of 50 parameters (**Figure 2B, right panel**), in general agreement with previously published results profiling fine locomotion in aged flies (of the Canton-S genetic background) with the flywalker (Cabrita *et al*., 2022), although the effects were slightly less pronounced in the current study and a few spatial parameter changes with age were in opposite directions in both studies. These differences could be due to the different genetic background or age of the animals profiled and compared in each study (7 vs 21 days in Cabrita e*t al.* (2022) compared to 2 vs 20 days in this study; **Figure 2B, left and right panels**). Further sampling under the same controlled conditions could help determine if these differences are reproducible and/or biologically significant.

Regardless of these differences, 20-day old *darhgef10[-/-]* females, exhibited a general worsening of their locomotor behavior as compared both to aged wild type controls (**Figure 2A, right column**) and to 2-day old *darhgef10[-/-]* female mutants, with the appearance of some significant changes not observed in control aged animals (**Figure 2B, right panel**). Few parameters were exclusively altered in aged *darhgef10[-/-]* females compared to control aged flies from this and the Cabrita *et al*. (2022) study, suggesting that most of the observed changes in aged *darhgef10[-/-]* females also occur in control aged flies, but to a weaker extent. An exception was the average area of all configurations, which was significantly increased in aged *darhgef10[-/-]* females but was either decreased or not-significantly affected in control aged flies (**Figure 2B**). This parameter reflects the area of all potential polygons formed by the legs during gaits involving four or more legs in stance phase. The increase in this measure indicates that aged *darhgef10[-/-]* females position their legs farther from the body, likely as a compensatory strategy to enhance stability during locomotion. Dimensionality reduction via principal component analyses further supported the significant difference between the walking behavior of aged *darhgef10* mutant females and both young mutants and aged control females (**Supplementary Figure S2**). The general aggravation of walking defects in aged *darhgef10* mutants relative to control aged animals indicated that dArhgef10 plays a significant role in mitigating the effects of aging on locomotor behavior in females. The appearance of phenotypes exclusively in aged *darhgef10* mutant females compared to aged controls and younger *darhgef10* mutants could further indicate specific requirements for *darhgef10* in aged animals.

To verify if *darhgef10* was also required for proper locomotor behavior in aged males, we assayed aged *darhgef10* hemizygous mutants (*darhgef10[-]/Y*) and controls (*darhgef10[+]/Y*) using the flywalker system. Compared to aged control males, aged *darhgef10[-]/Y* males exhibited very similar locomotor behavior changes to aged female mutants, with few exceptions, such as step frequency and tripod duration which were not significantly affected in aged mutant males, and a small significant increase in non-canonical index, which was not affected in aged mutant females (**Figure 2C, left column**). Hence, fine kinematic profiling using the flywalker showed that the *darhgef10* deletion significantly affected locomotor behavior in both female and male flies largely in similar ways, with a few sex-specific effects.

### Panneuronal *darhgef10* knockdown phenocopies walking kinematics defects of aged male *darhgef10* mutants

A mutation in human *ARHGEF10* identified in a family with subclinical slow-nerve-conduction velocity was associated with thin myelin sheaths (Verhoeven *et al.,* 2003), which are generated by Schwann cells, a type of glial cell of the peripheral nervous system. Wrapping glia, the equivalent axon-ensheathing cell in the *Drosophila* peripheral nervous system, is most similar to non-myelinating vertebrate Schwann cells, the “Remak” Schwann cells, which share a common developmental origin with the myelinating ones (Reed *et al.,* 2021), although wrapping glia also form myelin-like sheaths in the initial axonal segments of motor and sensory neurons of adults, performing therefore similar functions as both vertebrate Schwann cells (Rey *et al.,* 2023). Hence, we hypothesized that dArhgef10 activity would be required in glial cells for proper locomotion of aged flies. If this were correct, knockdown of *darhgef10* in glial cells of aged animals should phenocopy the locomotor phenotype of aged *darhgef10* mutants.

To test this, we used the flywalker system to analyze the locomotor activity of aged animals with RNA interference (RNAi)-mediated *darhgef10* knockdown (*>darhgef10-IR*) in glial cells using the pan-glial GAL4 driver transgenic line, *repo>* (*repo>darhgef10-IR*). Animals carrying either a copy of *repo>* or *>darhgef10-IR* alone served as controls for background and transgene insertion effects. As two control groups were analyzed, two data sets were generated using the flywalker system: one data set considered aged *repo>* males as the control group, and the other considered aged *>darhgef10-IR* males as the control group (**Supplementary Figure S3**). To facilitate the interpretation of the results, the columns that compare the glial *darhgef10* knockdown flies to each one of the control groups (*repo>darhgef10-IR* to *repo>* and *repo>darhgef10-IR* to *>darhgef10-IR* flies were integrated into a single column (**Figure 2C, middle column**). Flywalker analysis revealed statistically significant alterations only in three locomotor parameters in aged *repo*>*darhgef10*-IR males compared to both aged *repo*> and *>darhgef10*-IR control groups. Specifically, there was a slight reduction in swing speed variability (average standard deviation), a decrease in the non-canonical index, and a mild increase in the wave index. The reduced variability in swing speed suggests more consistent and coordinated leg movements. Furthermore, the lower non-canonical index indicates that aged *repo*>*darhgef10*-IR males perform fewer non-canonical gaits—unclassified yet stable gait patterns—suggesting improved locomotor coordination relative to both aged control groups. Importantly, out of all the locomotor parameters significantly altered in aged *darhgef10[-]/Y* mutants, only wave index was also significantly altered in the same direction in *repo>darhgef10-IR* males, suggesting a very limited effect of glial *darhgef10*-knockdown on male fly locomotor behavior and that this knockdown explains little or nothing of the *darhgef10[-]/Y* mutant locomotor phenotype.

As an alternative hypothesis, we verified if dArhgef10 activity was required in neurons, instead of glial cells, for proper maintenance of locomotion in aged males. For this, we carried out panneuronal *darhgef10* RNAi using the *57C10-GAL4* (*57C10>*) driver (Pfeiffer *et al.,* 2008) (*57C10>darhgef10-IR*) and assayed locomotion activity using the flywalker system. Controls consisted of *>darhgef10-IR* flies crossed to the “enhancer-less” GAL4 driver line, *pBDP>* (*pBDP>darhgef10-IR*) inserted at the same site (*attp2*) as *57C10>*. Flywalker results revealed that 43 out of 50 locomotor parameters were significantly altered in aged *57C10>darhgef10-IR* males in the same direction as in aged *darhgef10[-]/Y* males (**Figure 2C, right column**). These parameters included decreased speed, and increased swing speed, stance duration, duty factor, stance linearity, AEP footprint clustering, and average area of all 3-point contacts. Together, these results indicate that neuronal *darhgef10* is required for proper induced locomotor activity in aged *Drosophila* males, and that the *darhgef10*-dependent locomotor phenotype is better explained by the neuronal, rather than glial, *darhgef10* knockdown, contrary to our initial hypothesis.

### Spontaneous locomotor behavior is negatively affected by *darhgef10* deletion in aged females, but not aged males

The flywalker system provides high-resolution-tracking of walking kinematics typically in response to a brief airflow injected with an insect aspirator into the fly tunnel where flies are filmed. To investigate spontaneous locomotor behaviors such as initiation of movement and sleep, and also obtain an independent measure of coarse walking kinematics, such as speed and total distance covered and time spent walking, we tracked individual aged *darhgef10* flies and controls in open arenas for 2 h (**Supplementary Figure S4**). In contrast to the results obtained in the flywalker, we found no statistical difference between aged *darhgef10* mutant and control males in most spontaneous locomotor behaviors studied (**Figure 3**), except for a slightly reduced % pause time (pauses lasting less than 10 min), suggesting no or minimal effect of *darhgef10* deletion on male fly spontaneous locomotion. Aged mutant females, however, showed a statistically significant ∼50% decrease in speed and an increase in inactive time, which could be explained by an increase in pause time and sleep time, which per se are likely due to reduced walking bouts and increased sleeping bouts, respectively (**Figure 4**), compared to aged controls. These results indicated that female fly spontaneous locomotor behavior is strongly impaired by the *darhgef10* deletion. Altogether, these results suggested that *darhgef10* plays a much clearer role in the control of female spontaneous locomotor behaviors than that of males.

**Figure 3.**
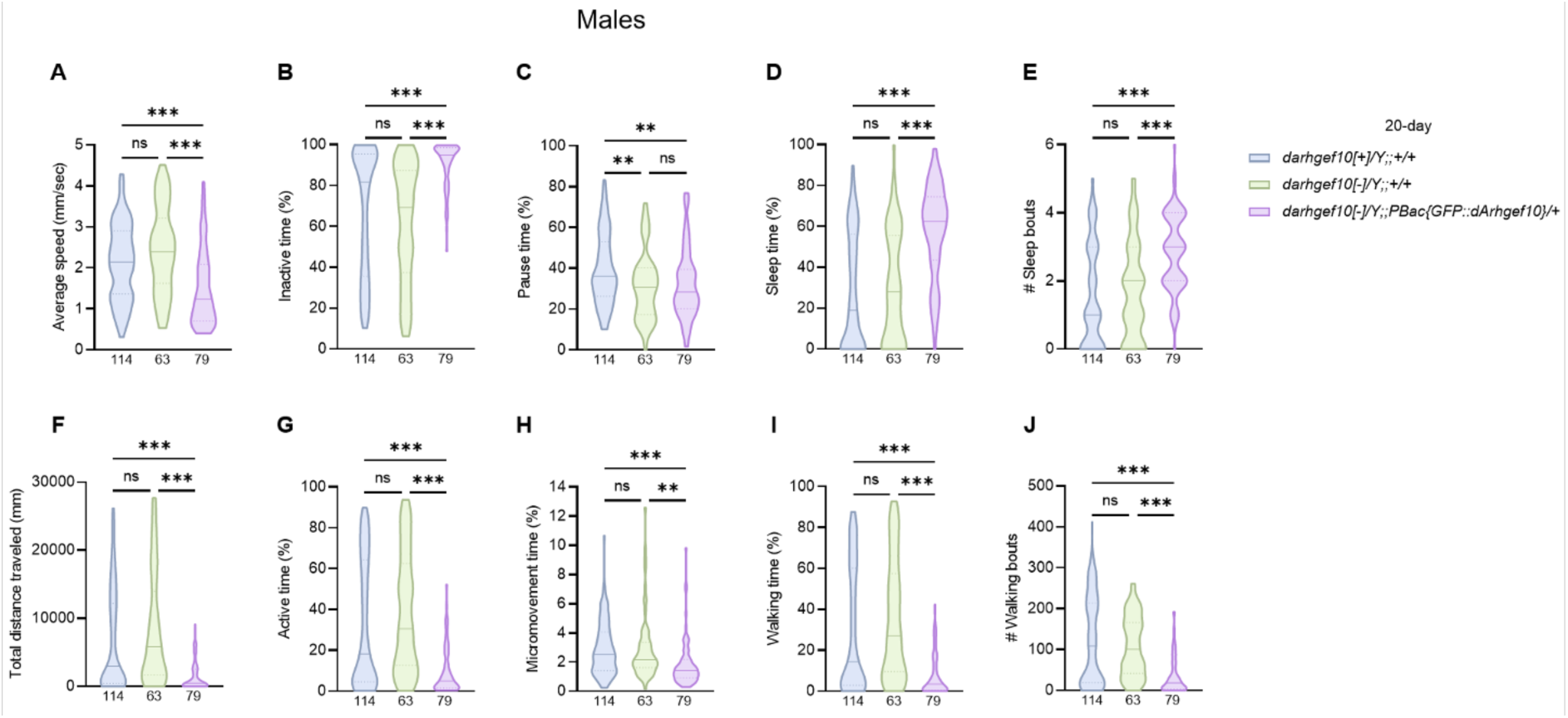
Spontaneous locomotor behavior is minimal or not affected by *darhgef10* deletion in aged males. Violin plots show locomotor parameters from the spontaneous locomotor behavior assay in wild-type controls (*darhgef10[+]/Y;;+/+*, blue), *darhgef10* knockouts (*darhgef10[-]/Y;;+/+*, green), and genetically rescued flies (*darhgef10[-]/Y;;PBac{GFP::dArhgef10}/+*, light violet) at 20 days post-eclosion. The central line indicates the median; lower and upper dashed lines indicate the 25th and 75th percentiles. Sample size (*N*) is shown at the base of each plot. Statistical significance was determined by one-way ANOVA followed by Tukey’s multiple comparisons test: *P* ≤ 0.05 (*), *P* ≤ 0.01 (**), *P* ≤ 0.001 (***), ns = not significant (*P* > 0.05).

**Figure 4.**
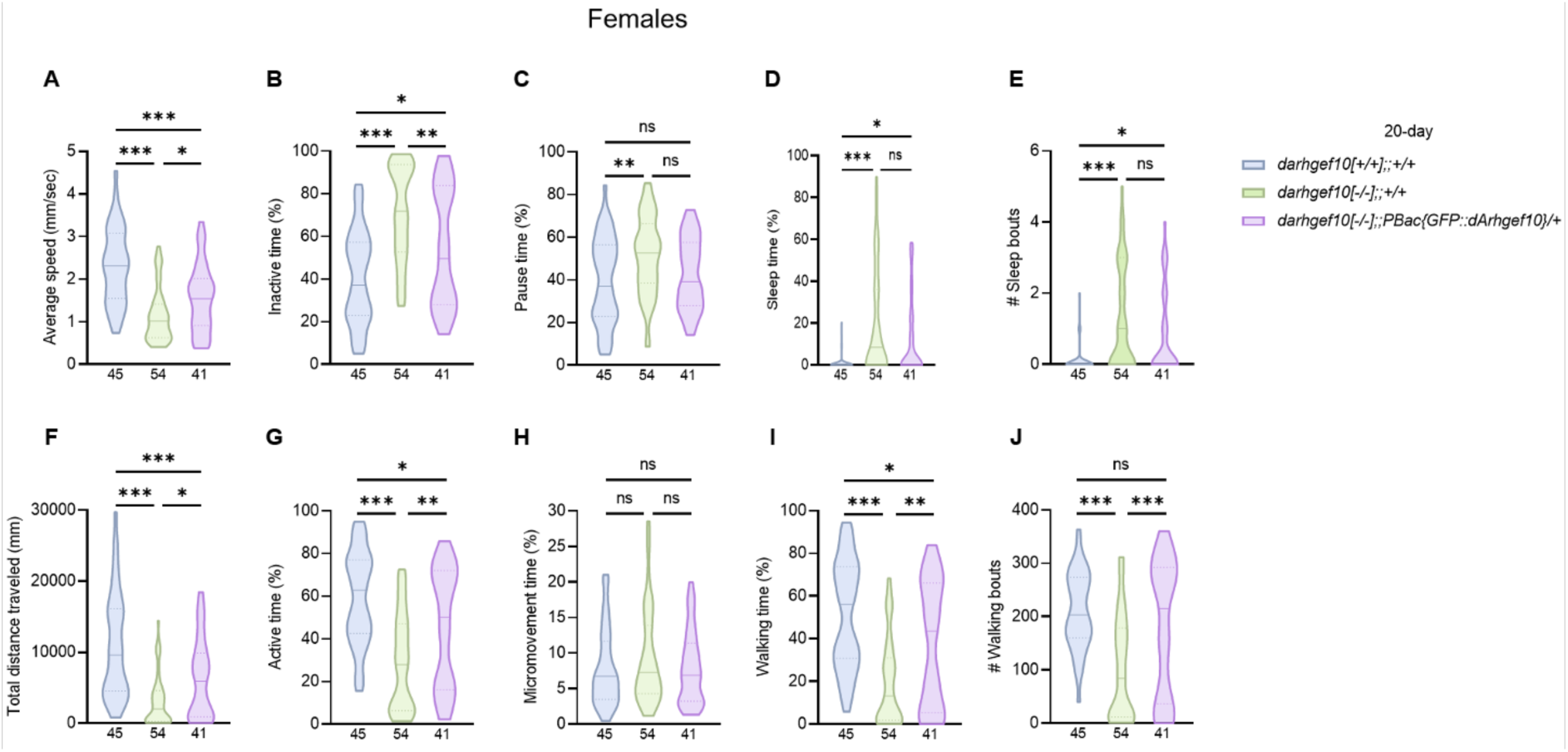
Spontaneous locomotor behavior is negatively affected by *darhgef10* deletion in aged females. Violin plots show locomotor parameters from the spontaneous locomotor behavior assay in wild-type controls (*darhgef10[+/+];;+/+*, blue), *darhgef10* knockouts (*darhgef10[+/+];;+/+*, green), and genetically rescued flies (*darhgef10[+/+];;PBac{GFP::dArhgef10}/+*, light violet) at 20 days post-eclosion. The central line indicates the median; lower and upper dashed lines indicate the 25th and 75th percentiles. Sample size (*N*) is shown at the base of each plot. Statistical significance was determined by one-way ANOVA followed by Tukey’s multiple comparisons test: *P* ≤ 0.05 (*), *P* ≤ 0.01 (**), *P* ≤ 0.001 (***), ns = not significant (*P* > 0.05).

### Sex-specific effects of a *GFP::darhgef10* rescue construct lacking isoforms *RC* and *RD*

To strengthen these claims genetically, we attempted to rescue *darhgef10[Df(1)ΔB]* with a *darhgef10* construct (*PBac{GFP::dArhgef10*}) located in the third chromosome (Di Pietro *et al.,* 2023), which consists of a carboxy (C)-terminally GFP-tagged dArhgef10 carrying most of the *darhgef10* locus except the promoters and other regulatory elements of isoforms C and D (**Figure 1B**) (see full genotype in **Supplementary Table 1** and *Drosophila* husbandry and stocks section from materials and methods for details). Unexpectedly, we found that the presence of the *GFP::darhgef10* rescue cassette in aged *darhgef10* mutant males negatively affected all locomotor parameters (**Figure 3A, B, D-J**), except pause time (**Figure 3C**), which coincidently was also the only parameter negatively affected by *darhgef10[Df(1)ΔB]* in aged males. In striking contrast, aged mutant females carrying the *GFP::dArhgef10* cassette showed improved spontaneous locomotor performance compared to aged *darhgef10* mutant females across all parameters that were affected by the mutation (**Figure 4**) even though the rescue did not reach statistical significance for pause and sleep time and the number of sleep bouts (**Figure 4C-E**). These results indicated that *GFP::dArhgef10* rescued the spontaneous locomotor effects of *darhgef10* mutation in females, but not in males, where it instead worsened spontaneous locomotor behaviors.

As *darhgef10* mutant males had major spontaneous locomotor behavior defects to begin with and the *GFP::darhgef10* construct provides only a subset of the total *darhgef10* transcripts, notably missing isoforms C and D–which encode long dArhgef10 protein isoforms with an extended N-terminal domain (**Figure 1B and C**)–, these findings indicate a sex-dependent difference in the activity of dArhgef10. One possibility is that males are particularly sensitive to an unbalanced set of available dArhgef10 isoforms. Males would require dArhgef10 isoforms C and/or D to regulate the effects of the other isoforms, so that in the absence of isoforms C and/or D, dArhgef10 activity becomes deleterious. On the other hand, the rescue of aged *darhgef10[Df(1)ΔB]* female locomotor behaviors by *GFP::darhgef10* argues that females do not require dArhgef10 isoforms C and D activity for proper locomotor behavior. To gain insight into these possibilities, we profiled *darhgef10* expression in control, mutant, and *GFP::darhgef10-*rescued *darhgef10* mutant male and female adults using quantitative reverse-transcriptase polymerase chain reaction (qRT-PCR) with isoform specific primers (**Figure 5A-F**). Results showed significant rescue of *darhgef10* mRNA levels by the *GFP::darhgef10* construct in both males and females (**Figure 5B-D**). Furthermore, we found clear male-specific enrichment of isoforms C and D, which were largely unaffected by *darhgef10[Df(1)ΔB]* or the *GFP::darhgef10* rescue construct, in contrast to the other isoforms (**Figure 5E-F**). As isoforms C and D coincidently are the isoforms that are not included in the *GFP::darhgef10* rescue construct, this is in line with our hypothesis that isoforms C and/or D are required for proper male locomotion, and are dispensable for female locomotion. Future work using targeted isoform rescue and RNAi could shed light into this.

**Figure 5.**
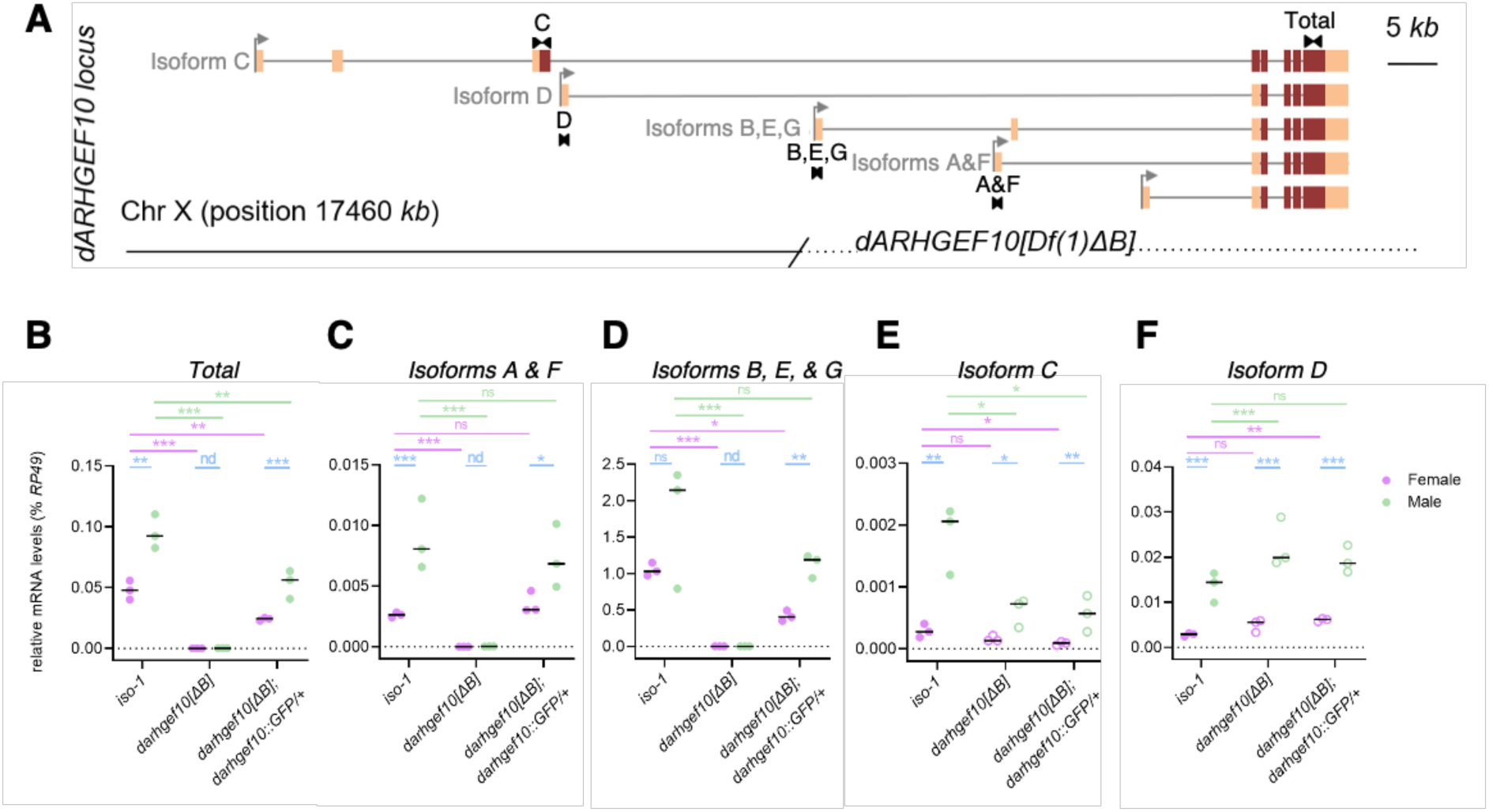
The isoforms C and D of *darhgef10* are enriched in male flies. **(A)** Eight *darhgef10* transcriptional isoforms are represented (“*A-G”*). Also depicted is part of the *darhgef10[Df(1)ΔB]* deletion, which covers the entire *darhgef10* coding region, including the DH and PH-like domain coding regions, for most isoforms except part of isoform C, which encodes a version of ARHGEF10 with an insect-specific N-terminal extension is not affected by the deletion. Primers used for isoform-specific and total *darhgef10* qRT-PCR analyses are depicted as black arrowheads. Isoform names are in gray. UTRs are in peach, and coding sequences are in brown. Transcriptional start sites are depicted as gray arrows. **(B-F)** Dot plots depicting the effect of sex, *darhgef10[Df(1)ΔB]* deletion, and *GFP::darhgef10-*rescue of *darhgef10[Df(1)ΔB]* on relative *darghef10* mRNA levels (depicted as % *Rp49,* housekeeping control) in 3-7 day-old male (green) and virgin females (magenta), as quantified by qRT-PCR using isoform specific primers for **(B)** total *darhgef10,* **(C)** isoforms *A & F*, **(D)** isoforms B, E, and G, **(E)** isoform C, and **(F)** isoform D. Controls (*iso-1*). N = 3 independent repeats. Horizontal bar, median. Empty dots in (E) and (F) depict the detection of the 5’ segments of the corresponding mutant transcripts. Statistical significance was determined by one-way ANOVA followed by Tukey’s multiple comparisons test: *P* ≤ 0.05 (*), *P* ≤ 0.01 (**), *P* ≤ 0.001 (***), ns = not significant (*P* > 0.05), nd, not done. Green, magenta, and blue lines and asterisks depict the statistical comparison between males of different genotypes, females of different genotypes, and males vs. females of the same genotype, respectively.

### *darhgef10* is required in neurons for spontaneous walking initiation and wakefulness in aged females

To investigate in which cells dArhgef10 may be required for the control of female spontaneous locomotor behavior, we assayed spontaneous locomotion of aged flies with *darhgef10* knockdown in glia, neurons, and muscle cells (three classes of cells involved directly or indirectly in locomotor control) using *repo>, neuronal synaptobrevin (nSyb)-GAL4* (*nSyb>*), and *myosing heavy chain (mhc)-GAL4* (*mhc>*), respectively. Aged male flies were also assayed as further controls, as we did not expect to find any effect of removing *darhgef10* in male tissues, as the germline knockout had virtually no effect on spontaneous locomotor behaviors (**Figures 3 and 4**). In general agreement with the profiling of induced walking kinematics, glial *darhgef10* knockdown had little effect on spontaneous behaviors, only slightly negatively affecting the average speed of spontaneous walking bouts (**Figure 6A, green**). Knockdown in muscle cells had no detectable effect on any spontaneous locomotor pattern (**Figure 6, blue**). In contrast and to our surprise, panneuronal *darhgef10* knockdown in males promoted more active time (**Figure 6G, pink**), which was explained by increased time spent walking and performing micromovements (**Figure 6H-I, pink**). The increased time walking could be explained by increased spontaneous walking bouts (**Figure 6J, pink**) rather than total distance traveled, which did not increase significantly. A separate experiment using another *nSyb* driver, *R57C10>* and another independent background control, confirmed these results (**Supplementary Figure S5**). The results above suggest that both glial and neuronal dArhgef10 make a relatively small, but significant contribution to proper spontaneous locomotor behaviors in aged males. Curiously, although *darhgef10* knockouts also spent slightly more time active and walking than controls, these differences were not statistically significant (**Figure 3G, I**), and time spent in micromovements were very similar to controls. Hence, although these results suggest that tissue-specific knockdown of *darhgef10* in males can reveal phenotypes that are not easily observable in *darhgef10* knockouts, they should be interpreted with caution and ultimately replicated with an independent RNAi line or *darhgef10* mutation.

**Figure 6.**
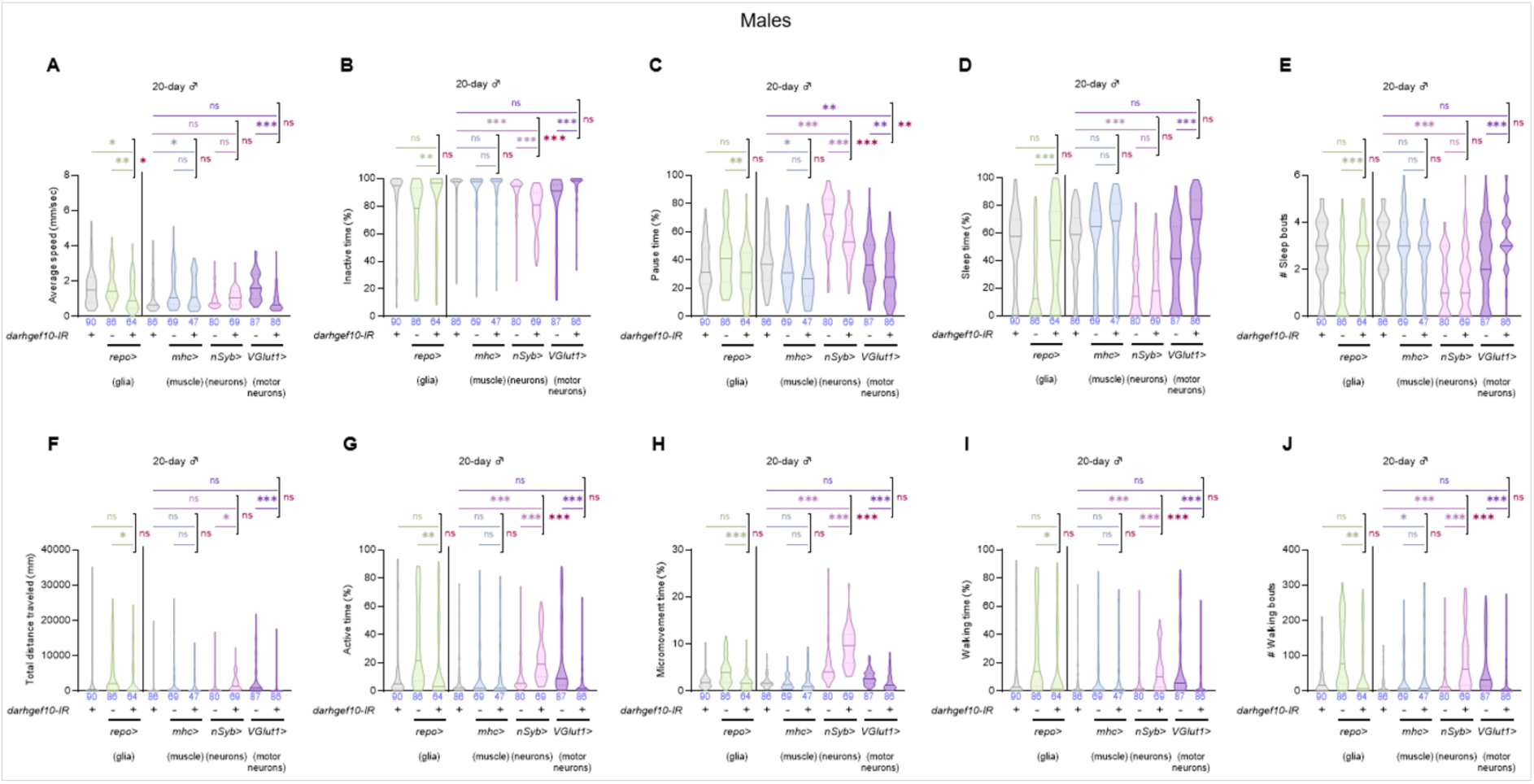
Tissue-specific *darhgef10* knockdown in males reveals phenotypes not observed in knockouts. Violin plots show locomotor parameters from the spontaneous locomotor behavior assay of 20-day-old males. Groups include controls (*>darhgef10-IR*, gray; *repo>*, green; *mhc>*, blue; *nSyb>*, pink; *VGlut1>*, violet) and tissue-specific knockdowns (*repo>darhgef10-IR*, green; *mhc>darhgef10-IR*, blue; *nSyb>darhgef10-IR*, pink; *VGlut1>darhgef10-IR*, violet). The black vertical line separates two independent experiments using distinct *>darhgef10-IR* control groups (gray) tested at 20 days post-eclosion. The central line indicates the median; lower and upper dashed lines indicate the 25th and 75th percentiles, respectively. Sample size (*N*) is shown at the base of each plot. Statistical significance was determined separately for each experiment using one-way ANOVA followed by Tukey’s multiple comparisons test: *P* ≤ 0.05 (*), *P* ≤ 0.01 (**), *P* ≤ 0.001 (***), ns = not significant (*P* > 0.05).

In aged females, however, the tissue-specific *darhgef10* knockdown results were generally consistent with those observed in aged female *darhgef10* mutants. Namely, glial and muscle knockdown had no consistent effect, except for a slight increase in micromovement time upon muscle *darhgef10* RNAi (**Figure 7H, blue**). In contrast, panneuronal *darhgef10* knockdown strongly affected spontaneous locomotion, significantly reducing average walking speed and active time compared to both controls (**Figure 7A, G, pink**). Aged female *nSyb>darhgef10* flies thus covered significantly less distance than their controls in the 2 h observation period (**Figure 7F, pink**). Animals also spent less time performing micromovements and walking (**Figure 7H-I, pink**), due to less walking bouts (**Figure 7J, pink**), and notably increased sleeping time, with ∼50% of the flies sleeping for >50% of the surveyed time (**Figure 7D, pink**). This extra time sleeping was due to an increased number of sleeping bouts (**Figure 7E, pink**). We conclude that there is strong support for a neuronal role for dArhgef10 in the control of spontaneous locomotor behaviors in aged females. The results above, taken together, also suggest there are sex-specific requirements for dArhgef10, most significantly in neurons, for the control of proper spontaneous locomotor behaviors in aged flies.

**Figure 7.**
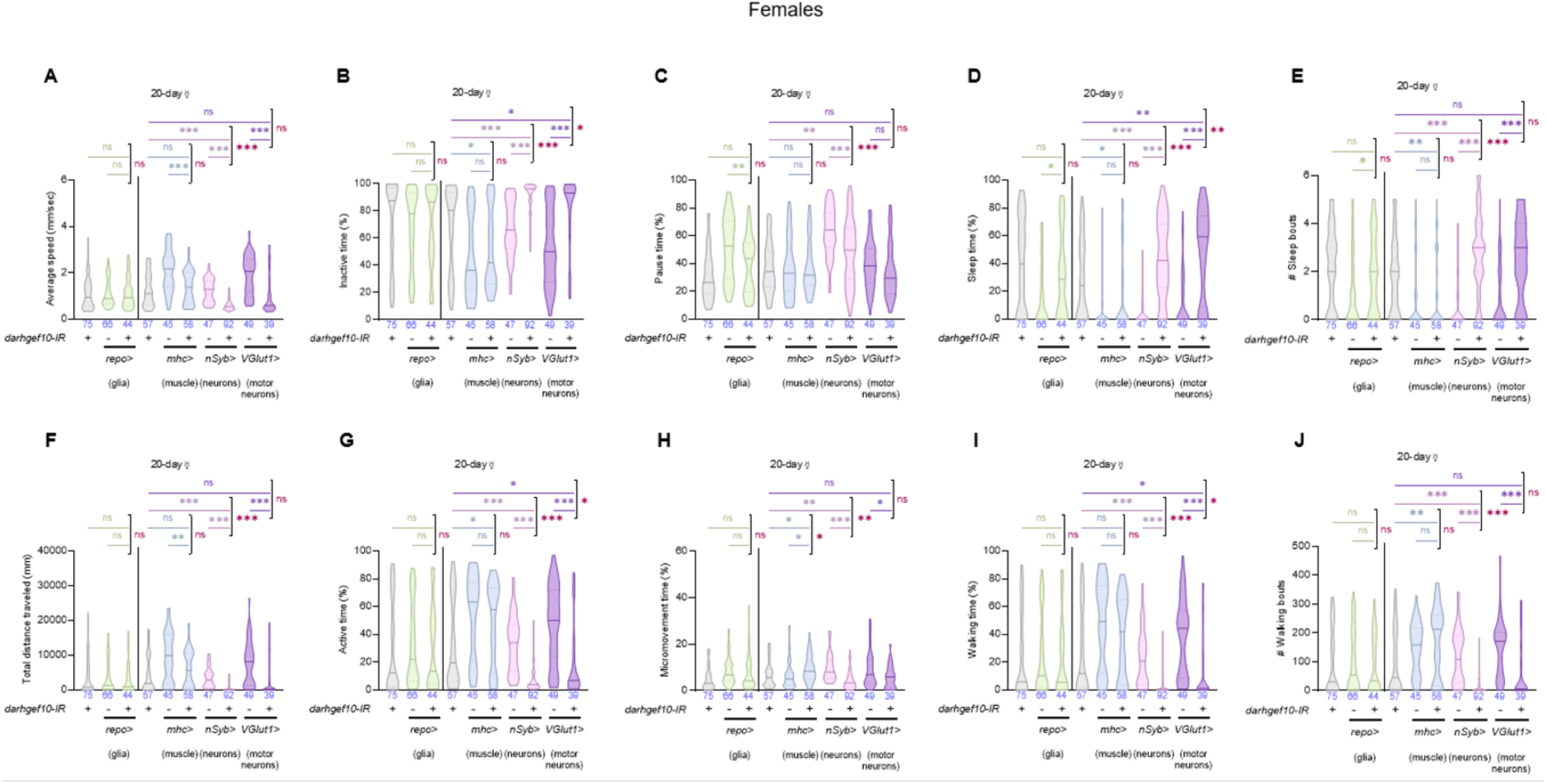
dArhgef10 is essential in neurons–more specifically in glutamatergic neurons–for proper spontaneous locomotion in aged females, suggesting a role in motor neurons. Violin plots show locomotor parameters from the spontaneous locomotor behavior assay of 20-day-old females. Groups include controls (*>darhgef10-IR*, gray; *repo>*, green; *mhc>*, blue; *nSyb>*, pink; *VGlut1>*, violet) and tissue-specific knockdowns (*repo>darhgef10-IR*, green; *mhc>darhgef10-IR*, blue; *nSyb>darhgef10-IR*, pink; *VGlut1>darhgef10-IR*, violet). The black vertical line separates two independent experiments using distinct *>darhgef10-IR* control groups (gray) tested at 20 days post-eclosion. The central line indicates the median; lower and upper dashed lines indicate the 25th and 75th percentiles, respectively. Sample size (*N*) is shown at the base of each plot. Statistical significance was determined separately for each experiment using one-way ANOVA followed by Tukey’s multiple comparisons test: *P* ≤ 0.05 (*), *P* ≤ 0.01 (**), *P* ≤ 0.001 (***), ns = not significant (*P* > 0.05).

### *darhgef10* is required in motor neurons for spontaneous walking initiation and wakefulness in aged females

We next hypothesized that dArhgef10 is required in motor neurons to regulate proper spontaneous locomotor behavior in aged flies. To test this, we knocked-down *darhgef10* using the *Vesicular glutamate transporter 1 (VGlut1)-GAL4* (*VGlut1>*) driver line, which is expressed in motor neurons, and tested fly spontaneous locomotion, as in the previous experiments. None of the effects seen in aged *nSyb>darhgef10-IR* males was phenocopied by *VGlut1>darhgef10-IR* (**Figure 6, violet**). This result could suggest that *darhgef10* is not required in neurons or in motor neurons of aged male flies for proper spontaneous locomotor behaviors, so that the neuronal effect observed with panneuronal RNAi experiments would be an experimental artifact. The fact that aged *darhgef10* knockout males do not present the same effects as aged males with panneuronal *darhgef10* RNAi could favor this interpretation. Alternatively, the absence of spontaneous locomotor behavior defects in aged *VGlut1>darhgef10-IR* males could suggest that dArhgef10 is required in other neuronal populations, rather than glutamatergic neurons, to reduce the amount of time the flies remain active (walking and performing micromovements) in the arenas. Further work is necessary to clarify to which extent dArhgef10 is critical for the control of spontaneous locomotor behaviors in aged males. However, it is safe to conclude that the contribution of dArhgef10 to coarse spontaneous locomotor behaviors in aged adult males is less clear and relatively smaller than its contribution to the fine induced walking kinematics (**Figure 2**).

In contrast to males, we observed that aged *VGlut1>darhgef10-IR* females more closely phenocopied aged *nSyb>darhgef10-IR* females, showing increased inactive time (**Figure 7B, violet**), which was mostly explained by a strong increase in sleep time (**Figure 7D, violet**). Aged *VGlut1>darhgef10-IR* females also showed decreased active time (**Figure 7G, violet**), which was characterized by less time spent walking (**Figure 7I, violet**). We conclude that *darhgef10* has a sexually dimorphic requirement in the nervous system for the execution of proper spontaneous locomotor behaviors, with the most striking finding being that glutamatergic neuron dArhgef10 promotes spontaneous activity and wakefulness in aged female flies, but not in aged male flies.

## DISCUSSION

Despite evidence that ARHGEF10 plays a role in the proper functioning of the nervous system and hence in the control of locomotor behavior in various organisms, including humans (Verhoeven *et al.,* 2003; Høyer *et al.,* 2014; Beutler *et al.,* 2014; Boora *et al.,* 2015; Ekenstedt *et al.,* 2014; Catusi *et al.,* 2021; Zhang *et al.,* 2022), the molecular and cellular mechanisms linking the gene to the behavior effects and to their age- and sex-specific effects are unclear. To gain insight into such mechanisms, we assayed the locomotor activity of *Drosophila melanogaster* flies lacking *darhgef10* with the hope to capitalize on the many advantages of this model for aging, behavior, and molecular genetics research.

We found that 20-day-old aged male and female locomotor activities were compromised to different extents in the absence of dArhgef10 activity. We used two different methods to study locomotion: the flywalker system, which assays fine induced walking kinematics, and free behavioral arena monitoring, which assays spontaneous locomotor behaviors, such as movement initiation and sleep. These systems calculate fundamentally distinct parameters, hence it is not completely surprising that the results we obtained did not always coincide across the assays. Most strikingly, fine walking kinematics of aged males was strongly affected by the loss of *darhgef10*, whereas these mutants performed fairly normally in the open behavioral arenas. In contrast, aged female mutant flies performed poorly in both assays. We interpret this as a clear indication that flies have sex-specific requirements for dArhgef10 to ensure proper behaviors, especially the spontaneous behaviors, being female flies the most sensitive to dArhgef10 levels. This sensitivity to dArhgef10 was not the only indication of sex-specific effects: a second strong indicative of sex-specific requirements was the rescue of aged *darhgef10* mutant female locomotor defects in the open behavioral arena by the *GFP::darhgef10* cassette, while aged *darhgef10* mutant males showed a clear worsening of their behavioral performance in the presence of this cassette. We hypothesize that this sex-specific response to the presence of GFP::dArhgef10 might be due to the incomplete representation of the panel of *darhgef10* isoforms in this rescue cassette. Namely, due to its fairly large size for a *Drosophila* gene, the Bacterial Artificial Chromosome (BAC) containing the *GFP::darhgef10* cassette encodes most isoforms, including those that are most highly expressed (isoforms B & E), but leaves out the two longest isoforms, isoforms C & D (**Figure 1B**) (Di Pietro *et al.,* 2023). Isoform D is a long transcript that encodes a similar protein as the other isoforms, but isoform C is predicted to encode a longer protein with an N-terminal extension (**Figure 1B and C**). Our qRT-PCR results suggested that this isoform is the one with sexual dimorphic expression: being upregulated 3-4-times in males, suggesting it has male-specific requirements (**Figure 5E-F**). Furthermore, as the *GFP::darhgef10* cassette can rescue locomotor defects of aged *darhgef10* mutant females, one would expect that it had no phenotype in aged *darhgef10* mutant males, which have no detectable phenotype in the spontaneous arenas to begin with. However, the cassette strongly negatively affects spontaneous locomotor behavior of aged *darhgef10* mutant males, suggesting that it could be acting as a gain-of-function in aged males. While we hypothesize that this is due to the absence of isoforms C & D, which seem to be more critical to males than females, at least for locomotion-Our findings could also be explained by a generally deregulated expression of *darhgef10* originating from the 3rd-chromosome-inserted *GFP::darhgef10* cassette, considering *darhgef10* is an otherwise X-linked gene. Our qRT-PCR analyses suggest this is not very likely, as the levels of *darhgef10* transcripts in rescued mutant male animal is similar to controls (with the exception of isoforms C and D) (**Figure 5B-F**). A possible caveat of these rescue experiments is the fact that the rescue construct includes three genes apart from the partial *darhgef10* locus, all of which are small and likely complete: *p-cup*, *CG8568*, and *CR44692*. It is therefore formally possible that any of these genes have sex-specific requirements in females. It is more difficult to explain their deleterious effect in males, but differential expression due to the ectopic location of the rescue construct could also play a role. Nevertheless, considering the general phenocopying of the mutant behavioral profiles by *darhgef10-*specific RNAi (**Figures 2, 6, and 7**), we strongly favor the interpretation that the rescue and deleterious effects of the cassette are exclusively due to *darhgef10*. The observations of sex-specific requirements for *darhgef10* in locomotor control is consistent with the sexually dimorphic displays and prevalences of symptoms in the central and peripheral diseases associated with *ARHGEF10*, such as schizophrenia (Li *et al.,* 2016), Autism Spectrum Disorder (ASD) (Napolitano *et al.,* 2022), and peripheral neuropathies (Hanewinckel *et al.,* 2016).

Previous research led us to hypothesize that dArhgef10 activity would be critical in glial cells for proper fly locomotion (Jungerius *et al.,* 2008). However, our results showed that glial-specific *darhgef10* knockdown was largely innocuous to locomotion, only weakly impairing the average spontaneous walking speed in open arenas, and this occurred only in aged males, not in females. This effect on speed also had weak support because aged *darhgef10* knockout males showed no clear defect in these assays, so it should be taken with extra caution. In contrast, panneuronal *darhgef10* RNAi strongly affected locomotor parameters in both sexes, although the effects differed in extent and characteristic. Specifically, panneuronal *darhgef10* RNAi and *darhgef10* mutation affected the fine induced walking kinematics of aged males to similar extents, but the same was not found for spontaneous behaviors such as movement initiation and sleep. Therefore we conclude that neuronal dArhgef10 is critical for aged male fine walking coordination, but it is largely dispensable for the other coarse behaviors observed in freely behaving aged animals in open arenas. While, as mentioned above, these assays quantify different parameters, walking speed was quantified in both assays, albeit in different ways. Nevertheless, it is difficult to understand why aged *darhgef10* mutant males did not show a consistent average speed reduction in the spontaneous behavior arenas, but did show reduced speed in the flywalker. One possibility is that aged male flies walk more quickly in response to stimuli, putting in an extra effort, which is more sensitive to dArhgef10 levels, than when they walk spontaneously in open arenas. Clearly, teasing and reconciling these apparently opposing results requires further work. For females, nevertheless, we observed strong evidence for a neuronal role for dArhgef10 in coordinating spontaneous locomotor behaviors in open arenas. We could also pinpoint this activity to motor neurons using *darhgef10* RNAi directed towards glutamatergic neurons. The most solid evidence–observed in *darhgef10* mutants, *nSyb>darhgef10-IR*, and *VGlut-1>darhgef10-IR* aged females–indicated that dArhgef10 is required in motor-neurons for proper spontaneous activity and wakefulness states, time spent walking and sleeping were decreased and increased, respectively, in all three conditions.

We conclude that *darhgef10* plays a role in fly locomotor behavior control, with different functions between males and females. Furthermore, the *darhgef10*-dependent locomotor phenotype is mainly explained by the absence of neuronal dArhgef10 activity. Besides that, the molecular and cellular mechanisms associated with the regulation of locomotion, in which *ARHGEF10* is involved, remain unknown.

## MATERIALS AND METHODS

### *Drosophila* husbandry and stocks

The following stocks were used in this study:

The *darhgef10* mutant (*darhgef10[-]*) is *darhgef10[Df(1)ΔB]; iso-2; iso-3,* and was generated in this study and is described below. The *darhgef10* background control (*darhgef10[+])* is *w[1118]; iso-2; iso-3* and was a gift from Luis Teixeira. *darhgef10[Df(1)ΔB]; +/+; PBac{GFP::darhgef10}/TM3* is *darhgef10[-];;PBac{GFP::darhgef10}/TM3* and was generated in our lab using a *GFP::darhgef10* rescue construct, which was a gift from Yohanns Bellaiche. *darhgef10[-];;PBac{GFP::darhgef10}/TM3* females were crossed with *darhgef10[+]/Y* males to obtain *darhgef10*-genetically-rescued males (*darhgef10[-]/Y;;PBac{GFP::dArhgef10}/+*), and *darhgef10[-]/Y;;PBac{GFP::darhgef10}/TM3* males were crossed with *darhgef10[-/-]* females to get *darhgef10*-genetically-rescued females (*darhgef10[-/-];;PBac{GFP::dArhgef10}/+*).

Fly stocks were maintained in standard cornmeal-agar medium at 20 °C. For the experiments animals were grown at 25 °C, and 70% humidity to achieve a controlled life cycle of about 10 days. All *D. melanogaster* stocks used in this work can be found in Table 2.1. The stocks were obtained either from the Bloomington Drosophila Stock Center (http://flystocks.bio.indiana.edu/), from other laboratories as a gift, or were generated in our laboratory, as depicted below.

**Table 2.1.**
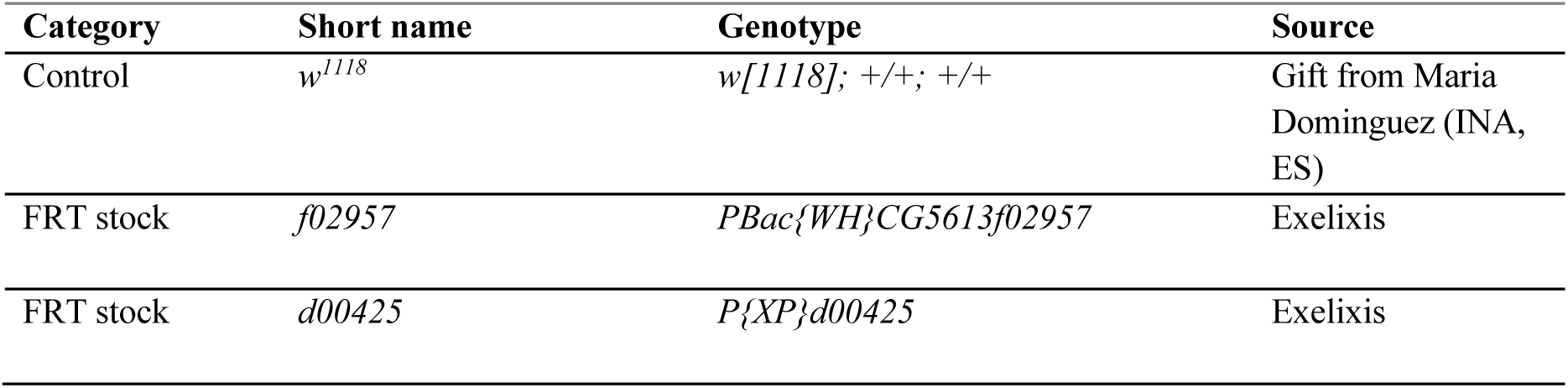
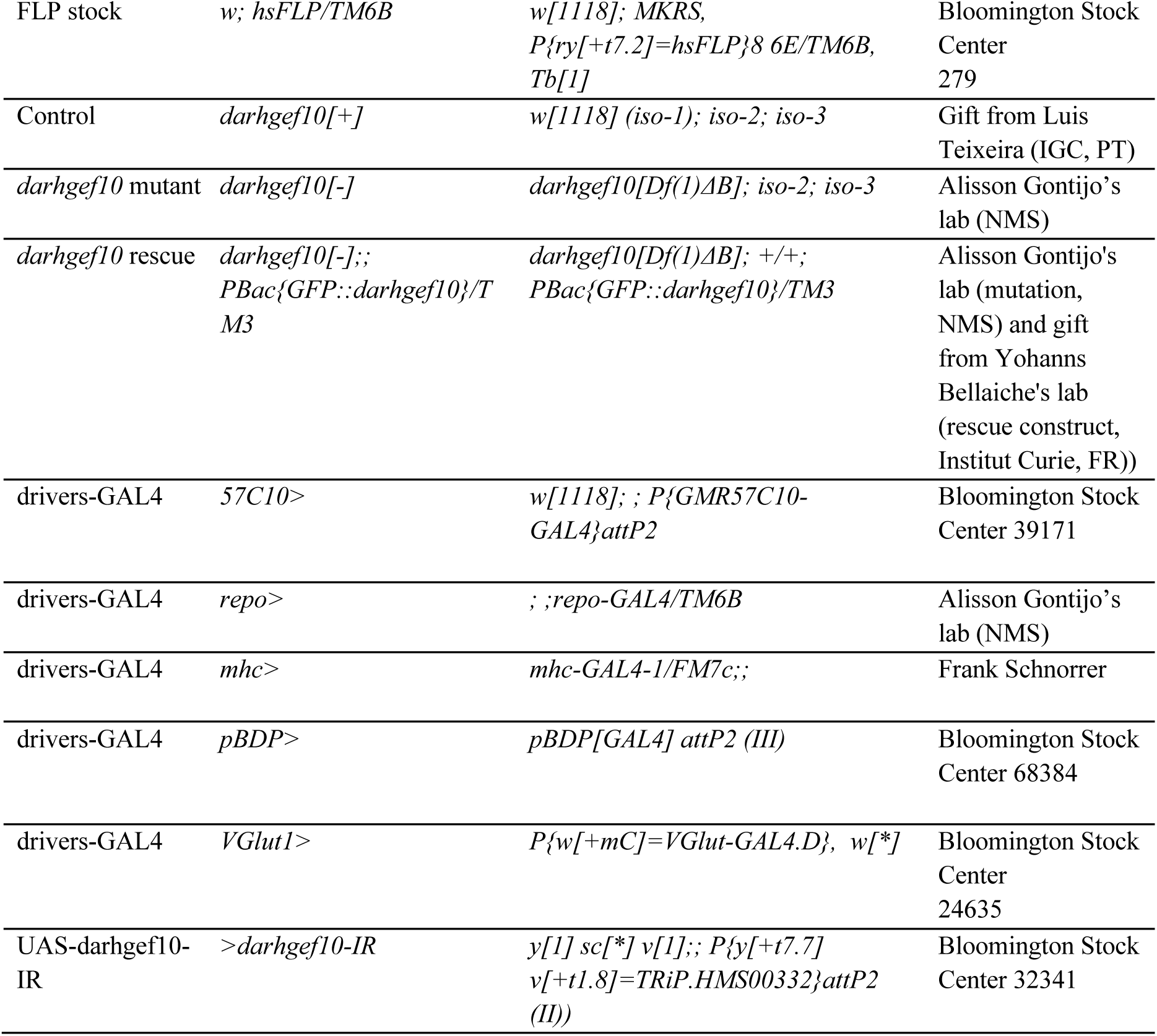
*D. melanogaster* stocks used in this work.

### Generation of the *darhgef10[Df(1)ΔB]* deletion

The *darhgef10[Df(1)ΔB]* mutant was generated by *FLP-FRT* deletion (Parks *et al.,* 2004). Flies carrying *FRT*-containing transposable elements flanking the region to be deleted (**Supplementary Figure S1A**) were selected: *f02957* (*WH*+) and *d00425* (*XP*-). Males carrying one of the *FRT* elements were mated to females carrying a FLP recombinase transgene (*w; hsFLP/TM6B*). From the progeny, males carrying both the transposable element and FLP recombinase sequences were selected and crossed to females carrying the second FRT element. Two days after, parents were removed from the vials and the progeny (containing both the FRT-bearing elements in trans and FLP recombinase) were subjected to daily 1-h heat-shocks for 5 consecutive days, by immersion of the vials into a 37°C water-bath. As *darhgef10* is located on the X chromosome, only virgin female flies were collected and mated to males containing balancer chromosomes (Binscinsy). Individual progeny (adult female *w*-virgin flies) were mated to Binsinscy males to generate additional progeny for PCR confirmation (using the the *WH+:XP-* hybrid element primers (WH5’_plus GACGCATGATTATCTTTTACGTGAC and XP5’_minus AATGATTCGCAGTGGAAGGCT) and analysis and to balance the stock in an isogenic background. Eventually, *Df(1)f02957-d00425* (*Df(1)ΔB*) hemi/homozygous mutant flies emerged and were incrossed to generate and maintain a stable homozygous mutant stock. The 2nd and 3rd chromosomes were substituted by those of the isogenic *w[1118]; iso-2; iso-3* line (Rubin and Lewis, 2000) with balancers, while backcrossing the X chromosome containing (*Df(1)ΔB*) 10 times.

The *Df(1)ΔB* deletion was confirmed by single-fly multiplex PCR (adapted from Carvalho *et al.,* 2009) using the primers *Rp49* (control) Fw: 5′-TTGAGAACGCAGGCGACCGT-3′ and Dm_Rp49_R Rv: 5’-AAGCCCAAGGGTATCGACAACA-3’ and the primers *darhgef10* (total) Fw: 5′-GGGTATAGCTGCCAATTCTGGAT-3 and Rv: 5′-ATTCGTGAAGGAGGACTACGATG-3′, which amplify a 288-bp and a 106-bp fragment, respectively in the presence of genomic DNA (gDNA) from controls (**Supplementary Figure S1B**).

### qRT-PCR

qRT-PCR was performed as previously described (Garelli *et al.,* 2015). Briefly, total RNA was isolated from male and female virgin adults aged 3-7 days using the Direct-zol RNA MiniPrep kit (Zymo Research) or NZY Total RNA isolation kit (NZYTech), following manufacturer’s instructions. Samples were macerated using pellet pestles, homogenized in 500 μl TRI Reagent or NZYol and centrifuged at 12000 g for 1 min, and treated with an extra DNAse treatment (Turbo DNA-free kit, Ambion, Life Technologies). cDNA synthesis was performed using the Maxima First Strand cDNA Synthesis Kit or NZY First-strand cDNA synthesis kit for RT–quantitative PCR (Thermo Scientific). qRT–PCR experiments were performed using Lightcycler 96 (Roche) or CFX96^TM^ (Bio-Rad) using the FastStart Essential DNA Green Master dye and polymerase (Roche). The final volume for each reaction was 10 μl, consisting of 5 μl of dye and polymerase (master mix), 2 μl of 10 × diluted cDNA sample and 3 μl of the specific primer pairs. Primers were designed using Primer BLAST or Primer3 and their efficiency was tested by serial dilution. ΔCq values (Cq*gene*−Cq*Rp49*) were used for statistical analyses. Data were analyzed by ANOVA followed by Tukey’s post-hoc tests using α = 0.05. In graphs, data were expressed as the geometric mean of %*Rp49* according to the formula: %*Rp49*=(2^−(Cq*gene*−Cq*Rp49*)) × 100 ± standard deviation of three biological repeats. The following primers were used:

*Rp49* (control)

Fw: 5′-TTGAGAACGCAGGCGACCGT-3′

Rv: 5′-CGTCTCCTCCAAGAAGCGCAAG-3′

*darhgef10* (total)

Fw: 5′-GGGTATAGCTGCCAATTCTGGAT-3′

Rv: 5′-ATTCGTGAAGGAGGACTACGATG-3′

*darhgef10* isoforms A & F

Fw: 5′-AGTTCTTTAGTGCGCCTTCATTGT-3′

Rv: 5′-GTTTCTTGGGCCTTTTCAGTGAGT-3′

*darhgef10* isoforms B, E & G

Fw: 5′-TTCTTTGGTTCGCACTTGTTGCT-3′

Rv: 5′-GCATTCGTGCCAAATCGTTGAGA-3′

*darhgef10* isoform C

Fw: 5′-GTATCCTGGCGCAGCTAAGTATC-3′

Rv: 5′-CACTGTATACGATGAGGCTGGAA-3′

*darhgef10* isoform D

Fw: 5′-GCACTCAATTCCAGGCCTAATTC-3′

Rv: 5′-TTCTCGTTTGCCCCGAAATACTA-3′

### Characterization of fly locomotor behavior

#### Flywalker system

The flies subjected to this experiment were grown in food vials under optimal development conditions (with a controlled number of eggs per vial to avoid overcrowding) to minimize any nutritional and/or environmental negative effect on their locomotor performance. Male adult animals were collected between 0-24 h after eclosion and kept in food vials for 20 days changed twice a week to ensure food supply and adequate media texture. The characterization of fly induced locomotor behavior was performed using 20-day-old flies as previously described by Mendes *et al.,* 2013. Briefly, flies were individually placed into a small chamber and filmed using a Photron (Tokyo, Japan) Mini UX-100 camera using a Nikon (Tokyo, Japan) AF 24-85 mm lens at 250 frames per second (fps). For each fly genotype, 10 animals were filmed twice to generate 20 videos. The videos obtained were analyzed using the flywalker software package written in MATLAB (Mendes *et al.,* 2013). Since this analysis generates errors in fly leg and body tracking, corrections had to be manually performed for each frame of a single video. From this analysis, a set of graphics (such as the stance traces, the step pattern, and the gait map) and a Microsoft Excel file containing all the locomotor parameters are generated for each fly (Mendes *et al.,* 2013).

#### Spontaneous locomotor behavior assay

This assay uses a locomotor activity monitoring device that consists of a support where a 3D-printed 30-well plate containing the flies, illuminated with LEDs, is placed, and a camera connected to a Raspberry Pi microcomputer for the acquisition of the videos (**Supplementary Figure S4A**). The 30-well plate consisted of 30 rectangular wells sized 20 x 10 mm at the top, 16 x 6 mm at the bottom, and 4 mm deep arranged in a 5 x 6 array (**Supplementary Figure S4B**).

Flies were anesthetized on an ice-cold surface, individually placed in the wells of the arena and covered with a transparent plastic adhesive tape. Usage of CO_2_ was avoided because it impairs spontaneous locomotor performance (Bartholomew *et al.,* 2015). After placing flies in the well plate, it was covered with a transparent plastic adhesive tape and placed in the locomotor activity monitoring device. Since the program used can detect the contour of each fly, creating a small yellow circle in the center of each fly’s contour (**Supplementary Figure S4B)**, their position can be mapped to a particular well. The (X, Y) coordinates of each yellow circle within the well are saved as the current position value and used to calculate the speed as the Euclidean difference with the position value of the fly detected in the same well in the preceding frame. The videos are acquired with a 1280 x 960-pixel resolution at 10 frames per second for 2 h and saved in 5-min-long fragments in .h264 format. After 2 h, recordings stop, and the well number, (X, Y) coordinates, speed, and time of each fly at each frame are saved in a .csv output file. The same file is subsequently processed to extract several spontaneous locomotor parameters: average speed, total distance traveled, active time, inactive time, micromovement time, walking time, pause time, sleep time, as well as the number of walking and sleep bouts. All parameters are computed individually for each fly based on a 2-h recording period. Average speed is calculated as the mean of all speed values greater than zero, while total distance corresponds to their sum (since speed reflects the distance covered per second). Active and inactive times are expressed as the percentage of speed values above or equal to zero, respectively. Active bouts are classified as micromovements if the distance covered is ≤50 px (6.76 mm), or as walking if >50 px. These are quantified as micromovement and walking times, representing the proportion of active bouts within each category. Inactive bouts are considered pauses if shorter than 600 s (10 min), or sleep if equal to or longer than this threshold; pause and sleep times reflect the percentage of inactive bouts in each category. To minimize circadian variability in locomotor activity (Gilestro *et al.,* 2012; Geissmann *et al.,* 2019), all 2-h recordings were performed during the same time window (11 a.m. to 3 p.m.).

### Data processing and statistical analysis

#### Flywalker system heatmap

Because locomotor parameters vary with speed (Mendes *et al.,* 2013; Mendes *et al.,* 2014; Cabrita *et al.,* 2022), all data were normalized by dividing each kinematic parameter by the corresponding fly’s speed. The mean value of each parameter was calculated and used to determine the effect size, expressed as the log₂ fold change (log₂ FC), defined as the logarithm base 2 of the ratio between the mean value of the experimental group and that of the control group. These log₂ FC values were visualized in a heatmap, where each column represents an experimental group compared to the control, and each row corresponds to a specific kinematic parameter measured using the FlyWalker software. In the heatmap, red shades indicate an increase in parameter value (positive log₂ FC), while blue shades indicate a decrease (negative log₂ FC), relative to control. White represents no change.

Statistical significance between control and experimental groups was assessed using a two-tailed t-test, with an alpha level of 0.05. Significance levels were indicated as follows: *P* ≤ 0.05 (*), *P* ≤ 0.01 (**), *P* ≤ 0.001 (***), and *ns* for non-significant values (*P* > 0.05). To constrain extreme values in the visualization, log₂ FC values greater than 1 were capped at 1, and values less than −1 were capped at −1.

In the case of the glial *darhgef10* knockdown experiment, two different control groups were used *(>darhgef10-IR* and *repo>*). Therefore, data from *repo>darhgef10-IR* flies were normalized to speed twice—once relative to each control group, as described above. Two sets of log₂ FC values were calculated: one for the comparison with *>darhgef10-IR* and another with *repo>* (**Supplementary Figure S3**). The final log₂ FC values used in the heatmap represent the average of these two comparisons. Thus, the middle column labeled “*repo>darhgef10-IR*” in Figure 2C integrates both analyses.

#### PCA

Principal component analysis (PCA) was performed as previously described (Cabrita *et al.,* 2022). Briefly, a regression model was generated using the control group to account for the effect of speed on locomotor parameters. Residual values for the experimental groups were then calculated based on this model and used as input for PCA.

All analyses were conducted using custom Python scripts (version 3.9.13, Anaconda distribution). The resulting PCA data were exported to Excel and visualized using GraphPad Prism. Box plots represent the median and the 25th and 75th percentiles (interquartile range, IQR). Whiskers extend to 1.5 times the IQR unless this range exceeds the data bounds—in such cases, the whiskers are limited to the minimum and maximum values. Values outside this range are considered outliers, according to Tukey’s method.

Statistical significance was assessed using Tukey’s multiple comparisons test when data followed a normal distribution, or the Kruskal–Wallis test followed by Dunn’s multiple comparisons test when the distribution was non-normal.

#### Spontaneous locomotor behavior assay data

The data generated by the program was plotted with GraphPad Prism in boxplots, and the statistical analysis was performed as outlined to the flywalker system data except for the assessment of statistical significance for comparisons between multiple conditions, in which the one-way ANOVA followed by Tukey’s multiple comparisons test were applied for normally distributed data.

## ACKNOWLEDGMENTS

We thank Luis Teixeira, Rita Teodoro, and Yohanns Bellaiche for stocks, and Rita Teodoro and members of the Integrative Biomedicine Laboratory for comments and discussion on the work. We thank Rafaela Pina for help in testing and setting up the open arenas. Stocks obtained from the Bloomington Drosophila Stock Center (NIH grant P40OD018537) were used in this study. TRiP RNAi stocks were used in this study (Office of the Director R24 OD030002). Research in the Integrative Biomedicine Laboratory was supported by the FCT (PTDC/BIA-BID/31071/2017; PTDC/MED-NEU/30753/2017 (LISBOA-01-0145-FEDER-030753, with financing from the Programa Operacional Lisboa 2020); EXPL/BIA-BID/1524/2021; EXPL/BIA-COM/1296/2021; 2022.03859.PTDC; 2022.08735.PTDC; 2023.15344.PEX), by the Research and Development Units iNOVA4Health (10.54499/UIDB/04462/2020) and Centre for Ecology, Evolution and Environmental Changes (cE3c) (10.54499/UIDB/00329/2020), by LS4FUTURE and CHANGE financed by the FCT/Ministério da Ciência, Tecnologia e Ensino Superior (Portugal), and by Congento LISBOA-01-0145-FEDER-022170, cofinanced by FCT/Lisboa2020; UID/Multi/04462/2019. Work in the Garelli lab was supported by Agencia Nacional de Promoción Científica y Tecnológica (PICT-2017-0254 and PICT-2020-01568), Consejo Nacional de Investigaciones Científicas y Técnicas (PIP11220150100182CO), and UNS-PGI-24/B288.ARMD. A.M.G. and F.H. were individually supported by Grants 10.54499/CEECINST/00102/2018/CP1567/CT0031 and SFRH/BPD/94112/2013 and 10.54499/DL57/2016/CP1457/CT0016, respectively. A.M., A.R.M.D. and M.G., were supported by the FCT fellowships, PD/BD/128445/2017, SFRH/BPD/94112/2013, and PD/BD/128003/2016, respectively. A.G. is a CONICET researcher. All reagents and fly strains generated in this study are available from the corresponding author without restriction.

## AUTHOR CONTRIBUTIONS

MGu, FH, AM, RZ, JM, ARMD, MGa, SP, AG, CM, and AMG performed research. MGu, FH, AM, CS, and AG obtained behavioral data. ARMD, SP, and AJ generated the KO. MGu, FH, ARMD, RZ, JM, and AMG performed molecular biology and qRT-PCRs. MGu, FH, AM, RZ, AG, CM, and AMG analyzed behavioral and expression data. AG established the open behavioral arena setup and wrote code. MGu and AMG wrote the manuscript with the help of all authors.

## BIBLIOGRAPHIC REFERENCES

Bartholomew, N. R., Burdett, J. M., VandenBrooks, J. M., Quinlan, M. C., & Call, G. B. (2015). Impaired climbing and flight behaviour in *Drosophila melanogaster* following carbon dioxide anaesthesia. Scientific reports, 5, 15298. 10.1038/srep15298

Beutler, A. S., Kulkarni, A. A., Kanwar, R., Klein, C. J., Therneau, T. M., Qin, R., Banck, M. S., Boora, G. K., Ruddy, K. J., Wu, Y., Smalley, R. L., Cunningham, J. M., Le-Lindqwister, N. A., Beyerlein, P., Schroth, G. P., Windebank, A. J., Züchner, S., & Loprinzi, C. L. (2014). Sequencing of Charcot-Marie-Tooth disease genes in a toxic polyneuropathy. Annals of neurology, 76(5), 727–737. 10.1002/ana.24265

Boora, G. K., Kulkarni, A. A., Kanwar, R., Beyerlein, P., Qin, R., Banck, M. S., Ruddy, K. J., Pleticha, J., Lynch, C. A., Behrens, R. J., Züchner, S., Loprinzi, C. L., & Beutler, A. S. (2015). Association of the Charcot-Marie-Tooth disease gene ARHGEF10 with paclitaxel induced peripheral neuropathy in NCCTG N08CA (Alliance). Journal of the neurological sciences, 357(1-2), 35–40. 10.1016/j.jns.2015.06.056

Cabrita, A., Medeiros, A. M., Pereira, T., Rodrigues, A. S., Kranendonk, M., & Mendes, C. S. (2022). Motor dysfunction in *Drosophila melanogaster* as a biomarker for developmental neurotoxicity. iScience, 25(7), 104541. 10.1016/j.isci.2022.104541

Carvalho, G. B., Ja, W. W., & Benzer, S. (2009). Non-lethal PCR genotyping of single Drosophila. BioTechniques, 46(4), 312–314. 10.2144/000113088

Catusi, I., Garzo, M., Capra, A. P., Briuglia, S., Baldo, C., Canevini, M. P., Cantone, R., Elia, F., Forzano, F., Galesi, O., Grosso, E., Malacarne, M., Peron, A., Romano, C., Saccani, M., Larizza, L., & Recalcati, M. P. (2021). 8p23.2-pter Microdeletions: Seven New Cases Narrowing the Candidate Region and Review of the Literature. Genes, 12(5), 652. 10.3390/genes12050652

Chaya, T., Shibata, S., Tokuhara, Y., Yamaguchi, W., Matsumoto, H., Kawahara, I., Kogo, M., Ohoka, Y., & Inagaki, S. (2011). Identification of a negative regulatory region for the exchange activity and characterization of T332I mutant of Rho guanine nucleotide exchange factor 10 (ARHGEF10). The Journal of biological chemistry, 286(34), 29511–29520. 10.1074/jbc.M111.236810

Di Pietro, F., Osswald, M., De Las Heras, J. M., Cristo, I., López-Gay, J., Wang, Z., Pelletier, S., Gaugué, I., Leroy, A., Martin, C., Morais-de-Sá, E., & Bellaïche, Y. (2023). Systematic analysis of RhoGEF/GAP localizations uncovers regulators of mechanosensing and junction formation during epithelial cell division. Current biology: CB, 33(5), 858–874.e7. 10.1016/j.cub.2023.01.028

Ekenstedt, K. J., Becker, D., Minor, K. M., Shelton, G. D., Patterson, E. E., Bley, T., Oevermann, A., Bilzer, T., Leeb, T., Drögemüller, C., & Mickelson, J. R. (2014). An ARHGEF10 deletion is highly associated with a juvenile-onset inherited polyneuropathy in Leonberger and Saint Bernard dogs. PLoS genetics, 10(10), e1004635. 10.1371/journal.pgen.1004635

Garelli, A., Heredia, F., Casimiro, A. P., Macedo, A., Nunes, C., Garcez, M., Dias, A. R. M., Volonte, Y. A., Uhlmann, T., Caparros, E., Koyama, T., & Gontijo, A. M. (2015). Dilp8 requires the neuronal relaxin receptor Lgr3 to couple growth to developmental timing. Nature communications, 6, 8732. 10.1038/ncomms9732

Geissmann, Q., Beckwith, E. J., & Gilestro, G. F. (2019). Most sleep does not serve a vital function: Evidence from *Drosophila melanogaster*. Science advances, 5(2), eaau9253. 10.1126/sciadv.aau9253

Gilestro G. F. (2012). Video tracking and analysis of sleep in Drosophila melanogaster. Nature protocols, 7(5), 995–1007. 10.1038/nprot.2012.041

Haga, R. B., & Ridley, A. J. (2016). Rho GTPases: Regulation and roles in cancer cell biology. Small GTPases, 7(4), 207–221. 10.1080/21541248.2016.1232583

Hanewinckel, R., van Oijen, M., Ikram, M. A., & van Doorn, P. A. (2016). The epidemiology and risk factors of chronic polyneuropathy. European journal of epidemiology, 31(1), 5–20. 10.1007/s10654-015-0094-6

Heasman, S. J., & Ridley, A. J. (2008). Mammalian Rho GTPases: new insights into their functions from in vivo studies. Nature reviews. Molecular cell biology, 9(9), 690–701. 10.1038/nrm2476

Hodge, R. G., & Ridley, A. J. (2016). Regulating Rho GTPases and their regulators. Nature reviews. Molecular cell biology, 17(8), 496–510. 10.1038/nrm.2016.67

Høyer, H., Braathen, G. J., Busk, Ø. L., Holla, Ø. L., Svendsen, M., Hilmarsen, H. T., Strand, L., Skjelbred, C. F., & Russell, M. B. (2014). Genetic diagnosis of Charcot-Marie-Tooth disease in a population by next-generation sequencing. BioMed research international, 2014, 210401. 10.1155/2014/210401

Jain, B. P., & Pandey, S. (2018). WD40 Repeat Proteins: Signalling Scaffold with Diverse Functions. The protein journal, 37(5), 391–406. 10.1007/s10930-018-9785-7

Joseph, J., Radulovich, N., Wang, T., Raghavan, V., Zhu, C. Q., & Tsao, M. S. (2020). Rho guanine nucleotide exchange factor ARHGEF10 is a putative tumor suppressor in pancreatic ductal adenocarcinoma. Oncogene, 39(2), 308–321. 10.1038/s41388-019-0985-1

Jungerius, B. J., Hoogendoorn, M. L., Bakker, S. C., Van’t Slot, R., Bardoel, A. F., Ophoff, R. A., Wijmenga, C., Kahn, R. S., & Sinke, R. J. (2008). An association screen of myelin-related genes implicates the chromosome 22q11 PIK4CA gene in schizophrenia. Molecular psychiatry, 13(11), 1060–1068. 10.1038/sj.mp.4002080

Khan, A., Ni, W., Lopez-Giraldez, F., Kluger, M. S., Pober, J. S., & Pierce, R. W. (2021). Tumor necrosis factor-induced ArhGEF10 selectively activates RhoB contributing to human microvascular endothelial cell tight junction disruption. FASEB journal: official publication of the Federation of American Societies for Experimental Biology, 35(6), e21627. 10.1096/fj.202002783RR

Li, R., Ma, X., Wang, G., Yang, J., & Wang, C. (2016). Why sex differences in schizophrenia?. Journal of translational neuroscience, 1(1), 37–42. https://www.ncbi.nlm.nih.gov/pmc/articles/PMC5688947

Lu, D. H., Liao, H. M., Chen, C. H., Tu, H. J., Liou, H. C., Gau, S. S., & Fu, W. M. (2018). Impairment of social behaviors in Arhgef10 knockout mice. Molecular autism, 9, 11. 10.1186/s13229-018-0197-5

Mendes, C. S., Bartos, I., Akay, T., Márka, S., & Mann, R. S. (2013). Quantification of gait parameters in freely walking wild type and sensory deprived Drosophila melanogaster. eLife, 2, e00231. 10.7554/eLife.00231

Mendes, C. S., Rajendren, S. V., Bartos, I., Márka, S., & Mann, R. S. (2014). Kinematic responses to changes in walking orientation and gravitational load in *Drosophila melanogaster*. PloS one, 9(10), e109204. 10.1371/journal.pone.0109204

Mohl, M., Winkler, S., Wieland, T., & Lutz, S. (2006). Gef10-the third member of a Rho-specific guanine nucleotide exchange factor subfamily with unusual protein architecture. Naunyn-Schmiedeberg’s archives of pharmacology, 373(5), 333–341. 10.1007/s00210-006-0083-0

Napolitano, A., Schiavi, S., La Rosa, P., Rossi-Espagnet, M. C., Petrillo, S., Bottino, F., Tagliente, E., Longo, D., Lupi, E., Casula, L., Valeri, G., Piemonte, F., Trezza, V., & Vicari, S. (2022). Sex Differences in Autism Spectrum Disorder: Diagnostic, Neurobiological, and Behavioral Features. Frontiers in psychiatry, 13, 889636. 10.3389/fpsyt.2022.889636

Parks, A. L., Cook, K. R., Belvin, M., Dompe, N. A., Fawcett, R., Huppert, K., Tan, L. R., Winter, C. G., Bogart, K. P., Deal, J. E., Deal-Herr, M. E., Grant, D., Marcinko, M., Miyazaki, W. Y., Robertson, S., Shaw, K. J., Tabios, M., Vysotskaia, V., Zhao, L., Andrade, R. S.,…Francis-Lang, H. L. (2004). Systematic generation of high-resolution deletion coverage of the *Drosophila melanogaster* genome. Nature genetics, 36(3), 288–292. 10.1038/ng1312

Pfeiffer, B. D., Jenett, A., Hammonds, A. S., Ngo, T. T., Misra, S., Murphy, C., Scully, A., Carlson, J. W., Wan, K. H., Laverty, T. R., Mungall, C., Svirskas, R., Kadonaga, J. T., Doe, C. Q., Eisen, M. B., Celniker, S. E., & Rubin, G. M. (2008). Tools for neuroanatomy and neurogenetics in Drosophila. Proceedings of the National Academy of Sciences of the United States of America, 105(28), 9715–9720. 10.1073/pnas.0803697105

Rubin, G. M., & Lewis, E. B. (2000). A brief history of Drosophila’s contributions to genome research. Science (New York, N.Y.), 287(5461), 2216–2218. 10.1126/science.287.5461.2216

Reed, C. B., Feltri, M. L., & Wilson, E. R. (2022). Peripheral glia diversity. Journal of anatomy, 241(5), 1219–1234. 10.1111/joa.13484

Rey, S., Ohm, H., Moschref, F., Zeuschner, D., Praetz, M., & Klämbt, C. (2023). Glial-dependent clustering of voltage-gated ion channels in *Drosophila* precedes myelin formation. eLife, 12, e85752. 10.7554/eLife.85752

Rümenapp, U., Freichel-Blomquist, A., Wittinghofer, B., Jakobs, K. H., & Wieland, T. (2002). A mammalian Rho-specific guanine-nucleotide exchange factor (p164-RhoGEF) without a pleckstrin homology domain. The Biochemical journal, 366(Pt 3), 721–728. 10.1042/BJ20020654

Shibata, S., Kawanai, T., Hara, T., Yamamoto, A., Chaya, T., Tokuhara, Y., Tsuji, C., Sakai, M., Tachibana, T., & Inagaki, S. (2016). ARHGEF10 directs the localization of Rab8 to Rab6-positive executive vesicles. Journal of cell science, 129(19), 3620–3634. 10.1242/jcs.186817

Verhoeven, K., De Jonghe, P., Van de Putte, T., Nelis, E., Zwijsen, A., Verpoorten, N., De Vriendt, E., Jacobs, A., Van Gerwen, V., Francis, A., Ceuterick, C., Huylebroeck, D., & Timmerman, V. (2003). Slowed conduction and thin myelination of peripheral nerves associated with mutant rho Guanine-nucleotide exchange factor 10. American journal of human genetics, 73(4), 926–932. 10.1086/378159

Winkler, S., Mohl, M., Wieland, T., & Lutz, S. (2005). GrinchGEF-A novel Rho-specific guanine nucleotide exchange factor. Biochemical and biophysical research communications, 335(4), 1280–1286. 10.1016/j.bbrc.2005.08.025

Zhang, Y., An, M. X., Gong, C., Li, Y. Y., Wang, Y. T., Lin, M., Li, R., & Tian, C. (2022). Single-copy Loss of Rho Guanine Nucleotide Exchange Factor 10 (*arhgef10*) Causes Locomotor Abnormalities in Zebrafish Larvae. Biomedical and environmental sciences: BES, 35(1), 35–44. 10.3967/bes2022.005

