## Supplementary Material for "Sex-specific control of locomotor behavior by neuronal Arhgef10 in aged *Drosophila*"

**This file contains:**
**Supplementary Tables S1-2.**
**Supplementary Figures S1-5.**

**Supplementary Tables**

**Supplementary Table S1. Full genotype of the animals used in Figures 2-6.**

| Short name | Genotype |
|---|---|
| >darhgef10-IR | w[1118]/y[1], sc[*], v[1]; +/+; P{y[+t7.7] v[+t1.8] =TRiP.HMS00332} attP2 (II) #32341 CG43658-IR darhgef10-IR/+ |
| repo> | w[1118]; +/+; repo-GAL4/+ |
| repo>darhgef10-IR | ; +/+; repo-GAL4/ P{y[+t7.7] v[+t1.8] =TRiP.HMS00332} attP2 (II) #32341 CG43658-IR darhgef10-IR |
| pBDP>darhgef10-IR | ; +/+; pBDP[GAL4] attP2 (III)/ P{y[+t7.7] v[+t1.8] =TRiP.HMS00332} attP2 (II) #32341 CG43658-IR darhgef10-IR |
| 57C10 >darhgef10-IR | w[1118]/y[1], sc[*], v[1]; +/+; P{GMR57C10-GAL4} attP2/P{y[+t7.7] v[+t1.8] =TRiP.HMS00332} attP2 (II) #32341 CG43658-IR darhgef10-IR |
| darhgef10[-];;PBac{GFP::darhgef10}/+ | darhgef10[Df(1)ΔB]; +/iso-2; PBac{GFP::dArhgef10}/iso-3 |
| mhc> | mhc-GAL4/w[1118]; +/+; +/+ |
| mhc>darhgef10-IR | mhc-GAL4/y[1], sc[*], v[1]; +/+; P{y[+t7.7] v[+t1.8] =TRiP.HMS00332} attP2 (II) #32341 CG43658-IR darhgef10-IR/+ |
| nSyb> | w[1118]; +/cyO ; nSyb-GAL4(III)/+ |
| nSyb>darhgef10-IR | y[1], sc[*], v[1]/w[1118]; +/cyO; nSyb-GAL4(III)/P{y[+t7.7] v[+t1.8] =TRiP.HMS00332} attP2 (II) #32341 CG43658-IR darhgef10-IR |
| VGlut1> | VGlut1-GAL4, w[*]/w[1118]; +/+; +/+ |
| VGlut1>darhgef10-IR | VGlut1-GAL4, w[*]/y[1], sc[*], v[1]; +/+; P{y[+t7.7] v[+t1.8] =TRiP.HMS00332} attP2 (II) #32341 CG43658-IR darhgef10-IR/+ |

**Supplementary Table S2. List and definition of the locomotor parameters generated by the flywalker system. b.u. - body units. Adapted from Mendes *et al.,* 2013, Mendes *et al.,* 2014, and Cabrita *et al.,* 2022.**

| Category | Locomotor parameter | Definition |
|---|---|---|
| **Step parameter** | Step Frequency (cycles/s) | Number of step cycles per second. |
| **Step parameter** | Step Period (ms) | Duration of one leg cycle consisting of one swing and one stance phase (Mendes *et al.,* 2013). |
| **Step parameter** | Swing speed (mm/s) | Speed of a leg in swing phase (Mendes *et al.,* 2013). |
| **Step parameter** | Step Length (μm) | Distance between two successive footprints of the same leg (Mendes *et al.,* 2013). |
| **Step parameter** | Swing Duration (s) | Duration of the swing phase (Mendes *et al.,* 2013). |
| **Step parameter** | Stance Duration (s) | Duration of the stance phase (Mendes *et al.,* 2013). |
| **Step parameter** | Duty Factor | Fraction of a step cycle in which a leg is on stance phase (Mendes *et al.,* 2014). |
| **Spatial parameter** | Footprint Alignment (μm) | Standard deviation from the average point of adjacent ipsilateral footprints projected onto the displacement axis (Mendes *et al.,* 2013; Mendes *et al.,* 2014). |
| **Spatial parameter** | Stance Linearity | Average difference between the stance traces generated by each leg during stance phase and a 5-point smoothed line (Mendes *et al.,* 2013; Mendes *et al.,* 2014). |
| **Spatial parameter** | AEP Footprint clustering (b.u.) | Standard deviation from the average position for all anterior extreme position (AEP) footprints. One AEP footprint corresponds to the position where the leg first contacts the glass after touchdown at the end of swing phase (Mendes *et al.,* 2013). |
| **Spatial parameter** | PEP Footprint clustering (b.u.) | Standard deviation from the average position for all posterior extreme position (PEP) footprints. One PEP footprint corresponds to the position of the leg at the end of the stance phase, just before the tarsus enters swing phase (Mendes *et al.,* 2013). |
| **Spatial parameter** | Stance Straightness | Ratio between the distance from AEP to PEP (displacement) and the path described by the tarsal contacts relative to the body (stance trace) (Cabrita *et al.,* 2022). |
| **Gait parameter** | Tripod Index | Percentage of frames in a video that display leg combinations defined by Tripod gait (Mendes *et al.,* 2013; Mendes *et al.,* 2014). |
| **Gait parameter** | Tetrapod Index | Percentage of frames in a video that display leg combinations defined by Tetrapod gait (Mendes *et al.,* 2013; Mendes *et al.,* 2014). |
| **Gait parameter** | Non-canonical Index | Percentage of frames in a video that display leg combinations defined by Non-canonical configuration (Mendes *et al.,* 2013; Mendes *et al.,* 2014). |
| **Gait parameter** | Wave Index | Percentage of frames in a video that display leg combinations defined by Wave gait (Mendes *et al.,* 2014). |

| | | |
|---|---|---|
| **Gait parameter** | Tripod Duration (ms) | Average of the duration of tripod gaits (Mendes *et al.,* 2014). |
| **Gait parameter** | Inter-tripod Transition Time (ms) | Duration of period between two tripod gaits (Mendes *et al.,* 2014). |
| **Stability parameter** | Body Displacement Ratio | Ratio between the distance walked and the path taken (Cabrita *et al.,* 2022). |
| **Stability parameter** | Average Area of all 3-point contacts | Area of the tripod support triangles. |
| **Stability parameter** | Average Area of all configurations | Area of the support polygons for all configurations. |

**Supplementary Figures**

**Supplementary Figure S1.**

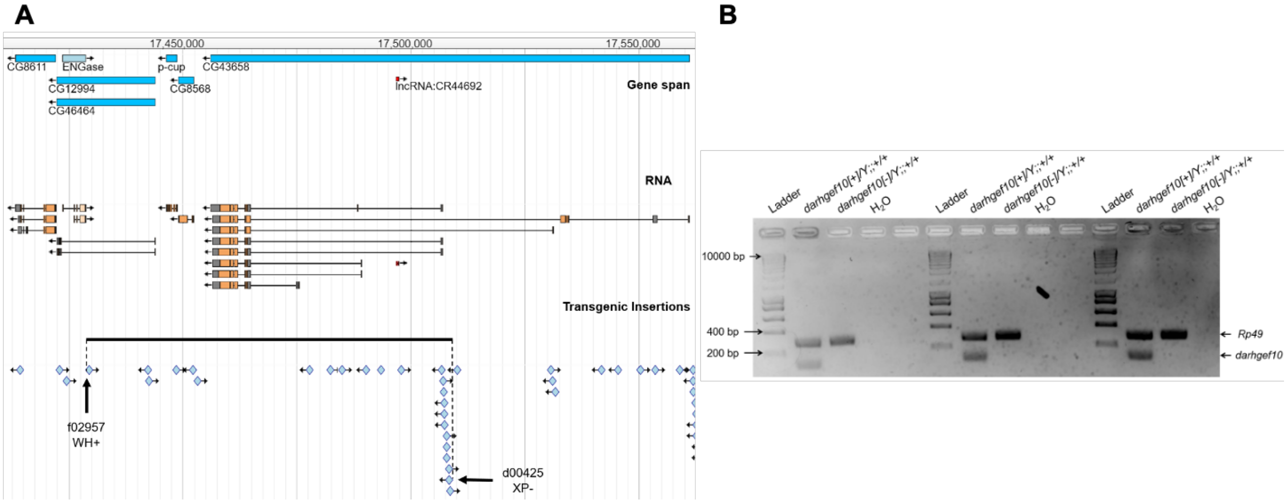


**Supplementary Figure S1. *darhgef10[Df(1)ΔB]* deletion. (A) JBrowse screenshot (from FlyBase,org) depicting the *dArhgef10* (*CG43658*) gene locus, eight *dArhgef10* transcripts (RNA), and the sites of mapped transgenic insertions in the *Drosophila melanogaster* genome.** Black arrows point to the transposons carrying the FRT sites that were used to generate the *darhgef10[Df(1)ΔB]* deletion (black line). **(B)** Multiplex PCR confirming the *darhgef10[Df(1)ΔB]* deletion. gDNA from one male control (*darhgef10[+]/Y;;+/+*) or *darhgef10[Df(1)ΔB]* mutant (*darhgef10[-]/Y;;+/+*)) was used in three independent multiplex PCR reactions using *Rp49* (control) and *darhgef10* primer pairs amplifying a 288- and 106-bp fragment, respectively. The *darhgef10* primers are located in the last common exon of the *arhgef10* gene, which is absent in animals carrying the *darhgef10[Df(1)ΔB]* deletion.

**Supplementary Figure S2.**

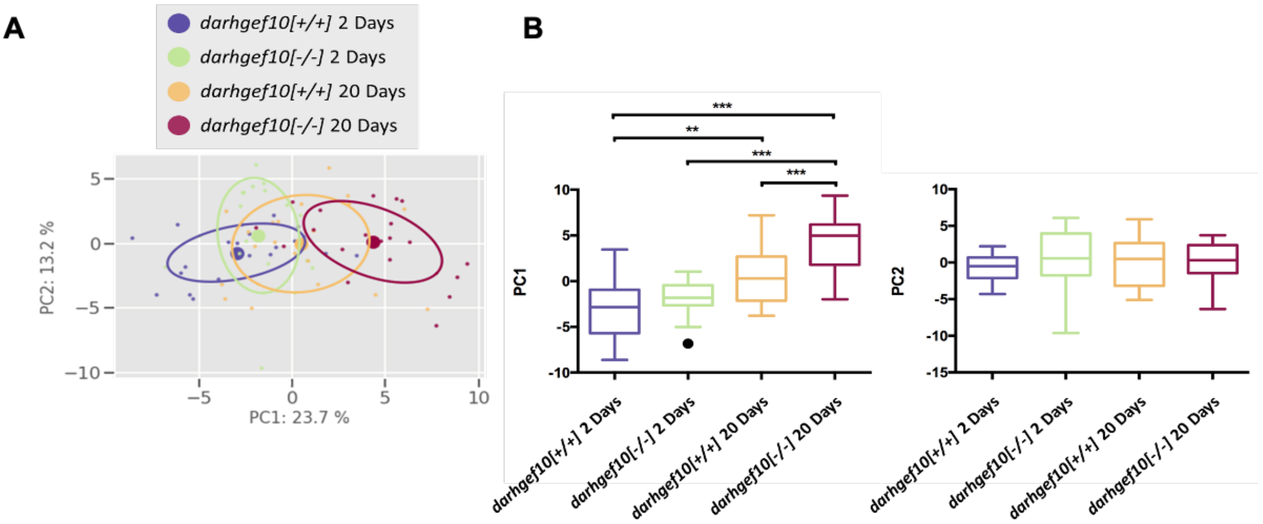


**Supplementary Figure S2. Aberrant and progressive locomotor defects in darhgef10 mutant females.** Principal Component Analysis (PCA) of all kinematic parameters. **(A)** 2D representation showing 50% of collected data (circles); small dots represent individual videos, and larger dots indicate the mean (n = 20 per condition). Component contributions are indicated on each axis. **(B)** Comparison of PC coordinates. Boxplots show medians (central line); lower and upper edges indicate the 25th and 75th percentiles; whiskers show the full range excluding outliers (black circles). Statistical significance was assessed using one-way ANOVA followed by Tukey's multiple comparisons test (for normally distributed data) or Kruskal–Wallis test followed by Dunn's multiple comparisons test (for non-normal data): $P \leq 0.05$ (*), $P \leq 0.01$ (**), $P \leq 0.001$ (***), ns = not significant ($P > 0.05$).

**Supplementary Figure S3.**

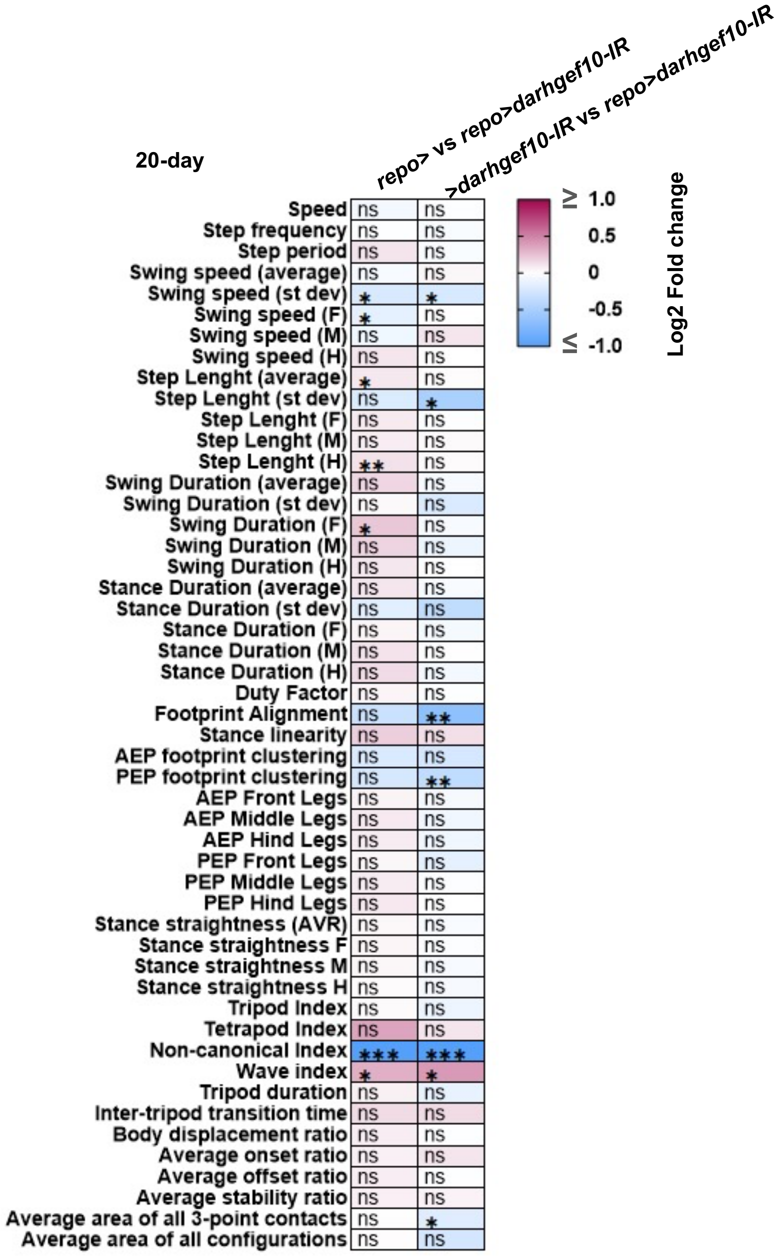


**Supplementary Figure S3. Analysis of the kinematic parameters of control and pan-glial *darhgef10* knockdown male flies, obtained through the Flywalker system.** Heatmap of the kinematic parameters of pan-glial *darhgef10* knockdown males (*repo>darhgef10-IR*) aged 20 days after eclosion. For each parameter values are matched to same age *repo>* (left column) and *>darhgef10-IR* (right column) control animals. The effect size ($Log_2$ Fold change) values are represented by a color code with red and blue shades indicating an increase or decrease relative to control, respectively. White indicates no variation. (n = 20 for each condition). Statistical significance was assessed by two-tailed t-test: $P \leq 0.05$ (*), $P \leq 0.01$ (**), $P \leq 0.001$ (***), ns = not significant ($P > 0.05$).

**Supplementary Figure S4.**

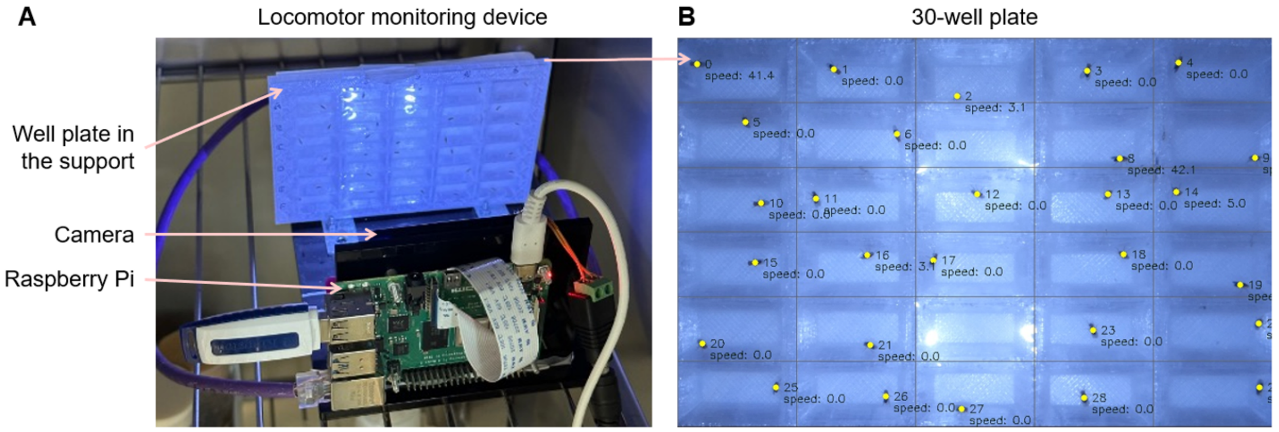


**Supplementary Figure S4. Spontaneous locomotor behavior assay. (A)** The locomotor monitoring device consists of a camera connected to the Raspberry Pi microcomputer and a support to place the 30-well plate. To assay fly locomotor activity, the 30-well plate containing the flies is placed in the support, and the fly locomotor behavior is recorded by a camera to obtain the videos. **(B)** Image of the well-plate containing the flies captured by the camera and processed by the program. The program can detect the position of each fly within each well (yellow circles) and calculate the speed of each fly. The top-left corner well is considered well 0 and the well on the bottom-right corner is considered well 29. Each fly is attributed the same ID number as the well.

**Supplementary Figure S5.**

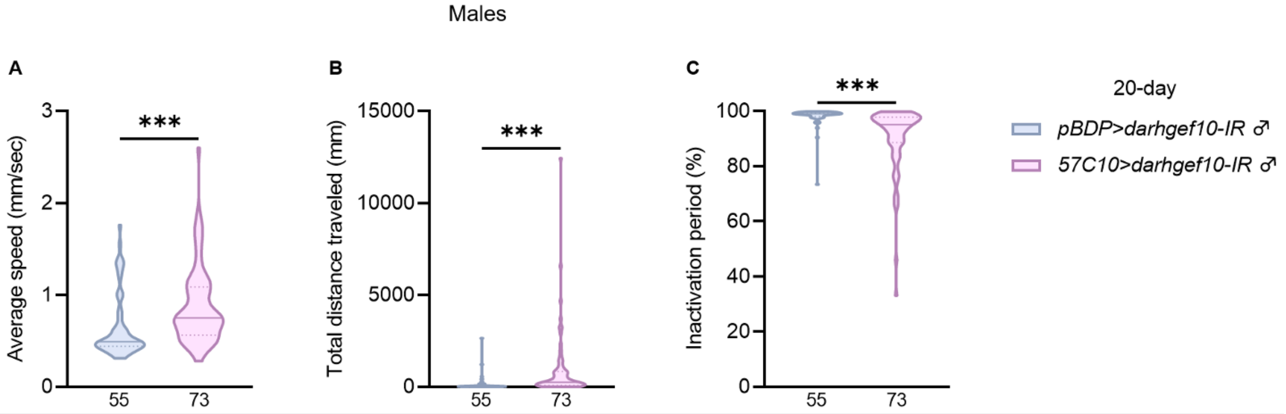


**Supplementary Figure S5. Panneuronal *darhgef10* knockdown improves induced locomotion in males.** Violin plots show spontaneous locomotor parameters for control (*pBDP>darhgef10-IR*, blue) and panneuronal knockdown (*57C10>darhgef10-IR*, pink) male flies at 20 days post-eclosion. Central lines indicate medians; dashed lines, 25th and 75th percentiles. *N* values are shown at the base of each plot. Statistical significance was assessed by Mann–Whitney test: $P \leq 0.05$ (*), $P \leq 0.01$ (**), $P \leq 0.001$ (***), ns = not significant ($P > 0.05$).
